# Epigenetic control of CD8^+^ T cell tissue homing and tissue resident memory T cell precursors by the histone methyltransferase SUV39H1

**DOI:** 10.1101/2024.05.13.593994

**Authors:** Guadalupe Suarez, Sandrine Heurtebise-Chrétien, Pierre-Emmanuel Bonté, Héloïse Beuchet, Jaime Fuentealba, Christel Goudot, Olivier Lantz, Sebastian Amigorena

**Author notes:** these authors contributed equally to the work (Sandrine Heurtebise-Chrétien, Pierre-Emmanuel Bonté).

## Abstract

Activation of CD8+ T cells leads to the differentiation of short-lived terminal effectors and memory precursors. Some of these memory precursors remain in lymphoid organs and become long-lived central memory T cells (T_CM_), while others home to non-lymphoid peripheral tissues early after antigen recognition and differentiate into tissue resident memory T cells (T_RM_). The early stages of memory precursor tissue homing and T_RM_ differentiation remain poorly understood. We show here that at steady state, during space-induced “homeostatic” expansion, and after flu infection, deletion of the histone 3-lysine 9 methyltransferase SUV39H1 in CD8^+^ T cells, increases the homing to non-lymphoid tissues (including liver, lungs, gut and skin). SUV39H1-defective cells in tissues express CD49d and differentiate into CD69+/CD103-T_RM_ after adoptive transfer or Flu infection. SUV39H1-defective T cells that accumulate in lungs are fully functional in both Flu re-infection and lung tumor models. We conclude that SUV39H1 restrains CD8^+^ T cell tissue homing and T_RM_ differentiation in WT mice. These results should encourage the use of SUV39H1-depletion in the context of adoptive T cell therapies to enhance tissue homing, thereby optimizing the efficiency of target cell eradication and long-term protection in the context of infection and cancer.

## Introduction

Effective immunological T cell memory is based on three fundamental features: 1) antigen mediated clonal expansion, which biases the memory T cell repertoire towards T cell receptors specific to pathogens encountered previously, 2) epigenetic landscape remodeling, that accelerates acquisition of effector functions in memory T cells, and 3) T cell re-distribution to peripheral tissues, that maximizes the chances of early pathogen re-encounter^1^. After infection or vaccination, antigen-specific T cells expand in lymphoid organs and differentiate into central memory T (T_CM_) and effectors (T_EFF_) cells. Some of these differentiating cells exit lymph nodes and home to non-lymphoid inflamed tissues, where they differentiate into tissue-resident memory T cells (T_RM_)^2^. Once in tissues, signals from the microenvironment contribute to inducing the expression of molecules essential for long-term tissue residency and survival, including CD69^3^ and CD103. T_RM_ also downregulate tissue egress receptors, such as S1PR1 and CCR7. Once established, T_RM_ provide local protection upon reinfection^4–6^ constitute a large proportion of the CD8^+^ T-cell memory population^7^ and are among the first to respond during secondary antigen encounters^5^. Lymphocyte homing to inflamed tissues involves a series of sequential migration steps requiring specific receptor-ligand interactions, including selectins, chemokines, and integrins. For example, the integrin VLA4 (a4b1, also known as CD49d/CD29) expressed by T cells binds to VCAM-1, mediating T cell arrest on endothelial cells before transmigration into the tissues. While it is know that T_RM_ precursors seed tissues during the effector phase of immune responses^8–12^, the precise nature of the T_RM_ precursors and their relationship to circulatory T cells, is still a matter of debate^1^.

The differentiation effector and memory T cell subtypes is accompanied by profound changes in the epigenetic landscape that maintain gene expression programs^13,14^. Studies on epigenetic changes during T_RM_ formation revealed both shared and tissue-specific epigenetic signatures^15^. Some of these changes occur early during immune responses, supporting the existence of poised T_RM_ precursors among tissue homing cells^16^. Post-translational modifications of histone N-terminal tails regulate many features of chromatin biology and gene expression and play crucial roles in cell development^17^, including in T cells^18–20^. Suppression of position effect variegation 3-9 1 (SUV39H1) was the first mammalian histone lysine methyltransferase (KMT) identified^21^. Together with its mammalian paralog SUV39H2, SUV39H1 introduces the second and third methyl groups on lysin 9 of histone H3 tails (H3K9me2 and H3K9me3). Through its KMT activity, SUV39H1 plays essential roles in heterochromatin formation and spreading^22–24^ and is one of the most conserved epigenetic silencing systems, even more than DNA methylation^24^. H3K9me3 is important for silencing lineage-inappropriate genes^25^ and for establishing linage cell identity during embryogenesis^26^, as well as an impediment to differentiated cells reprograming^27^. Previous work from our laboratory showed that H3K9me3 and SUV39H1 silence Th1-related genes during CD4^+^ T cell commitment to Th2 cells^18^. More recently, we showed that during effector CD8^+^ T cell differentiation, SUV39H1-mediated silencing of stemness genes is necessary to promote the establishment of cell identity, leading to the accumulation T_CM_ cells in lymphoid tissues^19^. Finally, we also showed that in tumor models, including CAR T cells, SUV39H1 defect leads to a dual phenotype, increased T_CM_ in lymphoid organs and increased effectors in tissues^28,29^.

Here, we analyze the role of SUV39H1 in T cell tissue homing and residence. We show SUV39H1 ablation in CD8^+^ T cells promotes increased homing to non-lymphoid tissues, T_RM_ precursor differentiation and CD69^+^ CD103^-^ T_RM_ cells accumulation at steady state, upon space-induced “homeostatic” expansion, and after flu infection. We conclude that in WT CD8^+^ T cells, SUV39H1 restrains tissue homing and T_RM_ differentiation. Together with our previous studies showing that SUV39H1 ablation enhances central memory and long-term T cell persistence, these results should encourage the use of SUV39H1 depletion or inhibition to enhance T cell-mediated protection in the context of adoptive T cell therapies and vaccination.

## Results

### SUV39H1-deficiency leads to the accumulation of CD8^+^ tissue-homing CD8^+^ T cells at steady-state

We have previously shown that SUV39H1 silences stemness genes during effector differentiation of CD8^+^ T cells upon *Listeria monocytogenes* (*Lm*) infection^19^. During *Lm* infection in SUV39H1 KO mice, not only *Lm*-specific T_CM_ accumulate in the spleen, but also CD69^+^ CD8^+^ T cells accumulate in the liver. These results suggest that SUV39H1 deficiency, in addition to promoting T_CM_ formation, may affect the differentiation of T_RM_ cells. To investigate this possibility, we first quantified CD8^+^ T cells in tissues at steady state in SUV39H1-deficient mice and WT littermates. As shown before, T_CM_ (CD62L^+^ CD44^+^) in lymphoid organs, including spleen and inguinal lymph nodes (iLN) accumulated among SUV39H1-deficient CD8^+^ T cells (Supp Fig 1 A, ^19^). CD62L^-^ CD44^+^ CD8^+^, but not CD4^+^, T cells also accumulated in non-lymphoid organs, including the liver (Supp Fig 1 A). We observed a similar increased accumulation of CD62L^-^ CD44^+^ CD8^+^ T cells in tissues of conditional SUV39H1^flox/flox^ CD4-Cre mice (Cre+) mice compared to WT (Cre-) littermates (Fig 1 A and Supp Fig 1 B). In contrast, these cells did not accumulate in the small intestines of SUV39H1^flox/flox^ CD4-Cre+ mice, where the great majority of CD8^+^ T cells were CD62L^-^ CD44^+^ (Fig 1 A).

**Figure 1:**
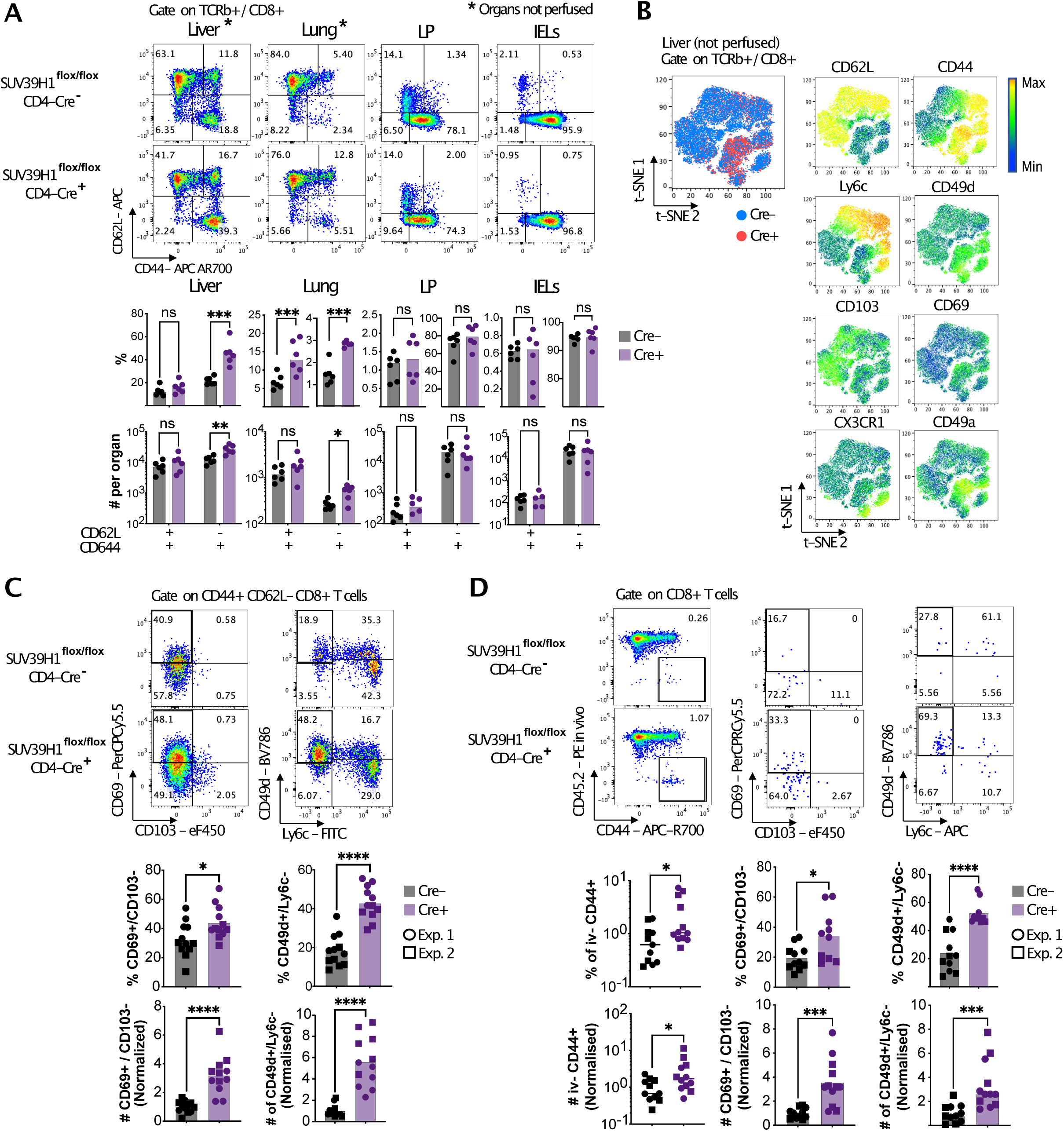
SUV39H1-deficiency leads to the accumulation of CD8^+^ tissue-homing CD8^+^ T cells at steady-state. **A.** Representative dot plots and quantification of CD44 and CD62L expression in CD8^+^ T cells from the indicated non lymphoid organs in conditional SUV39H1 KO (Cre+) or WT (Cre-) littermates. **B.** t-SNE plots of merged flow cytometry data gated on total CD8^+^ T cells from the liver of 5 conditional SUV39H1 KO (Cre+) and 5 WT littermates (Cre-) at steady state. See Supp Table 1. **C.** Representative dot plots and quantification of CD69^+^ CD103^-^ and CD49d^+^ Ly6c^-^ populations in CD44^+^ CD8^+^ T cells from liver of conditional SUV39H1 KO mice (Cre+) or WT littermates (Cre-) at steady state. **D.** Representative dot plots and quantification of intravascular staining negative (iv-) cells among lung CD8^+^ T cells and CD69^+^ CD103^-^ or CD49d^+^ Ly6c^-^ populations in iv-CD44^+^ CD8^+^ T cells from conditional SUV39H1 KO mice (Cre+) or WT littermates (Cre-) at steady state. Each dot represents a mouse, different shapes represent different experiments. * p<0.05 ; ** p<0.01 ; *** p< 0.001 ; **** p<0.0001 ; ns: not significant by multiple unpaired t test with Holm-Šídák correction (**A** and **B**) or Mann Whitney test (**C** and **D**).

SUV39H1 KO mice did not show any defect in thymic T cell development (Supp Fig 1 C) and the number of naïve CD8^+^ T cells in lymphoid organs were like those in WT mice (Supp Fig 1 D), suggesting normal T cell development. Accumulation of CD62L^-^ CD44^+^ cells in the liver was not dependent on the age of the animals (Supp Fig 1 E), excluding any possible effect of early aging on the accumulation of CD8^+^ T cells in tissues of SUV39H1-defficient mice^7^.

Unsupervised analysis of expression of memory and resident memory markers by flow cytometry (Supp Table 1) reveals that SUV39H1-deficient cells express higher levels of the T_RM_ marker CD69 and the integrins CD49d and CD49a, while expression of the circulatory marker Ly6c and the terminal effector marker CX3CR1 were low (Fig 1 B). Proportions and numbers of both CD69^+^ CD103^-^ and CD49d^+^ Ly6c^-^ CD8+ T cells were increased in livers of conditional SUV39H1-deficient mice as compared to WT littermates (Fig 1 C). To further validate tissue homing of these cells, as opposed to blood contaminating cells, we stained cells intravascularly (iv) with anti-CD45.2 mAbs for 5’ before mice euthanasia^30^. The extravascular (iv-) fraction of lung CD8^+^ T cells was increased in conditional SUV39H1 KO mice, both in percentage and cell numbers per organ (Fig 1 D). Moreover, the iv-fraction from SUV39H1-defficient mice was enriched in CD69^+^ CD103^-^ and CD49d^+^ Ly6c^-^ cells like in the liver. Altogether these results show that lack of SUV39H1 has a T cell intrinsic effect on CD8^+^ T cells accumulation in non-lymphoid tissues.

To analyze more in detail these tissue-associated CD8^+^ T cells, we next purified CD69^+^ Ly6c^-^ T_RM_ from the liver of both WT (Cre-) and conditional SUV39H1-defficient mice (Cre+) and used RNAseq and TCRseq to analyze gene expression and TCR clonality. Both SUV39H1 sufficient and deficient CD8^+^ T cells displayed high levels of computed T_RM_ signatures, showing no differences when comparing signature enrichment between Cre- and Cre+ cells, (Supp Fig 1 F), and showed very similar list of upregulated genes compared to naïve CD8^+^ T cells (Supp Fig 1 G). TCR-beta sequencing analysis of liver CD8^+^ CD69^+^ Ly6c^-^ T cells showed no increase in clonality among SUV39H1-deficient cells (compared to WT cells, Supp Fig 1 H), indicating that accumulation of these cells in tissues is not due to increased clonal size. Altogether our results show that SUV39H1-deficiency leads to the accumulation of T_RM_-like CD8^+^ CD69^+^ CD103^-^ cells at steady state.

### SUV39H1 restrains naïve CD8^+^ T cell differentiation into tissue homing cells

To evaluate the intrinsic role of SUV39H1 in CD8^+^ T cell tissue homing, we adoptively transferred naïve WT and SUV39H1-deficient OT-I CD8^+^ T cells at a 1:1 ratio into lymphopenic RAG-KO mice (Fig 2 A). Transferring naïve CD8^+^ T cells into lymphopenic hosts induces their expansion and promotes differentiation of T_RM_ cells in the gut after 27 days^31^. As shown in Fig 2B, between 33 and 35 days after transfer, SUV39H1-defficient cells accumulate up to 10 times more than WT cells in non-lymphoid organs. The KO:WT ratio was slightly over 1 in lymphoid organs, indicating a clear bias of SUV39H1-defective T cells for homing to non-lymphoid tissues, compared to lymphoid organs (Fig 2 B). Like at steady state, SUV39H1-deficient OT-I cells accumulated more than WT cells in the iv-compartment of the lung upon space-induced “homeostatic” proliferation (Fig 2 B, C and D). Both percentages and numbers of CD49d^+^ Ly6c^-^ cells were higher among SUV39H1-deficient OT-I cells in all tested organs (Fig 2 D). Like at steady state, CD49d^+^ Ly6c^-^ cells in iv-lung compartments expressed high levels of CD69 both in WT and KO cells (Fig 2 F). These results show that SUV39H1 restrains CD8^+^ naïve T cells differentiation into non-lymphoid tissue homing, CD49d-expressing, effector and/or memory cells in a cell intrinsic manner.

**Figure 2:**
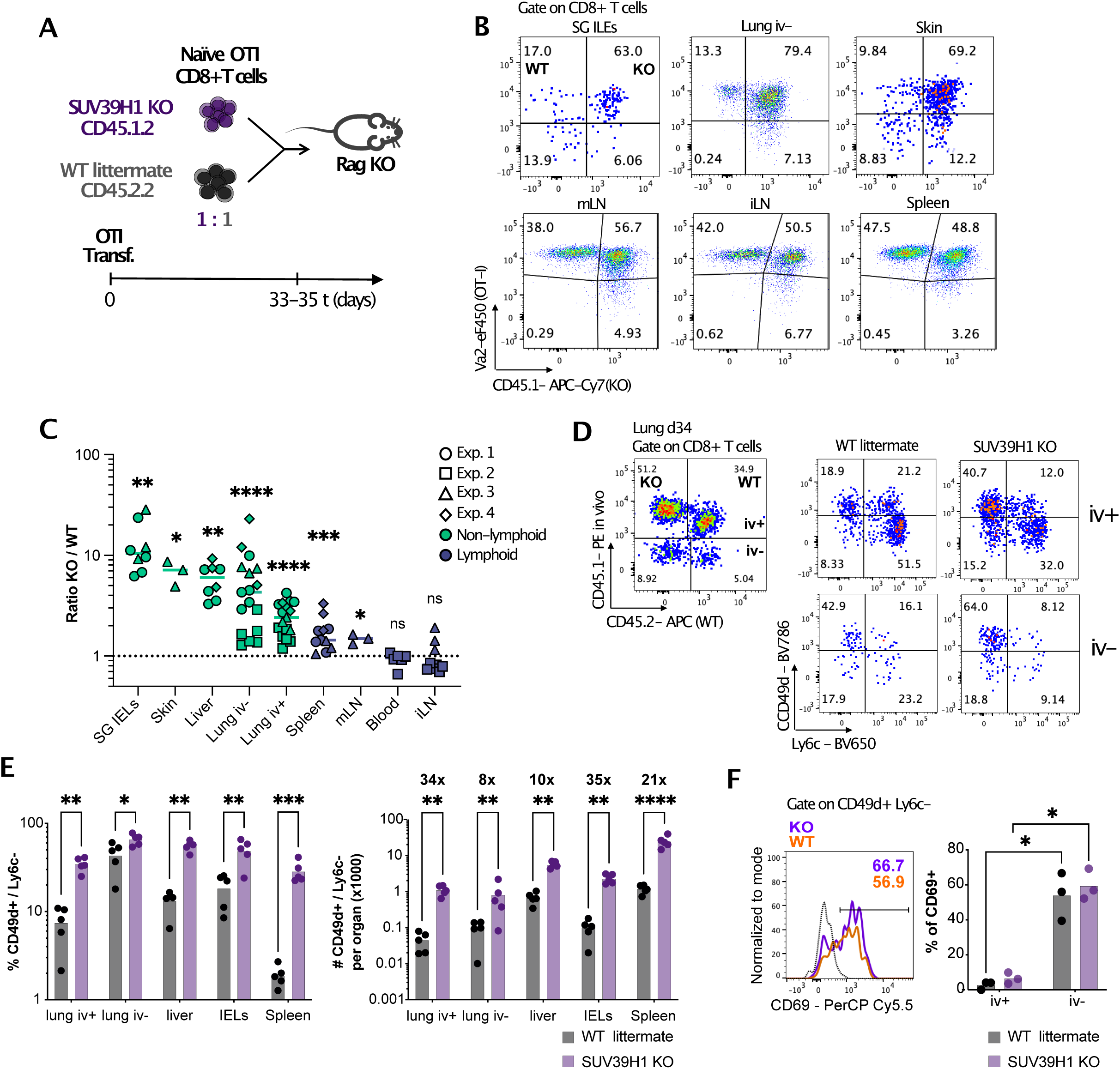
SUV39H1 restrains naïve CD8^+^ T cell differentiation into tissue homing cells. **A.** Scheme of the experiment. **B.** Representative dot plots of WT (CD45.1-) and SUV39H1 KO (CD45.1+) OTI cells (Vα2+) in the indicated organs 33 days after adoptive transfer. **C.** Quantification of the ratio between SUV39H1 KO and WT OTI cells within each organ. **D.** Representative dot plot of intravascular (iv) staining (**left**) and expression of CD49d and Ly6c (**right**) in each population at day 34 post transfer in CD8^+^ T cells from lungs from Exp. 2 in **C**. **E.** Quantification of the percentage and absolute numbers of CD49d^+^ Ly6c^-^ cells among SUV39H1KO or WT OTI cells in the indicated organs from Exp 1 in **C**. Numbers represent the fold increase between KO and WT condition for each organ. **F.** Histogram of CD69 expression in iv-CD49d^+^ Ly6c^-^ CD8^+^ T cells and proportions of CD69+ cells among iv+ and iv-CD49d^+^ Ly6c^-^ CD8^+^ T cells in lungs from Exp. 4 in **C**. Numbers represent p values. Each dot represents a mouse, different shapes represent different experiments. * p<0.05 ; ** p<0.01 ; *** p< 0.001 ; **** p<0.0001 ; ns: not significant by one sample t test compared to 1 (**C**), multiple ratio paired t test with Holm-Šídák correction (**E**) and multiple paired t test (**F**).

### CD8^+^ T_RM_ differentiation upon flu infection is enhanced in the absence of SUV39H1

We next investigated the role of SUV39H1 in CD8^+^ tissue-homing and T_RM_ formation after flu infection, a well-stablished model of T_RM_ differentiation in vivo^32–34^. In lungs of flu-infected mice, the extravascular (iv-) proportion of CD8^+^ T cells was increased as compared to uninfected mice (Supp Fig 2 B). No differences were observed between SUV39H1-deficient and sufficient T cells. Among these flu-induced lung-homing cells, however, we observed an accumulation of CD69^+^ CD103^-^ CD8^+^ T cells in SUV39H1-deficient mice, 35 days after PR8 flu infection (Supp Fig 2 B).

To evaluate the functionality of the SUV39H1 KO T_RM_ cells in lungs after flu infection, we made use of an heterosubtypic flu infection model in which protection to secondary infection is dependent on CD8^+^ T_RM_ cells^33,34^ (Fig 3 A). Mice were rechallenged 28 days after the first infection to avoid waning of the T_RM_ population in the lungs, which depends on the replenishment by circulating T_EM_ cells^34^. We observed an accumulation of CD49d^+^ Ly6c^-^ among the extravascular (iv-) CD8^+^ T cells in the lungs of conditional SUV39H1 KO (Cre-) mice, 28 days after X31 infection (Fig 3 B and Supp Fig 2 C and D). These cells also express high levels of CD127 and CXCR3, and low levels of KLRG1, suggesting that they constitute true memory, rather than long-lived effector populations (Supp Fig 2 E). Three days after the secondary infection with PR8 flu, we observed an overall accumulation of CD8^+^ T cells in the lung tissue (iv-), and an increase in the numbers of CD49d^+^ Ly6c^-^ and CD69^+^ CD103^-^ CD8^+^ T cells among CD44^+^ iv-cells (Fig 3 C). Finally, we also observed relatively lower viral loads in the lungs of SUV39H1 conditional KO mice 3 days after PR8 secondary infection, compared to WT littermates in mice that were previously infected with X31 flu strain (Supp Fig 2 F). Altogether these results suggest that SUV39H1 prevents the accumulation of protective CD49d^+^ CD69^+^ CD8^+^ T_RM_ cells upon flu infection.

**Figure 3:**
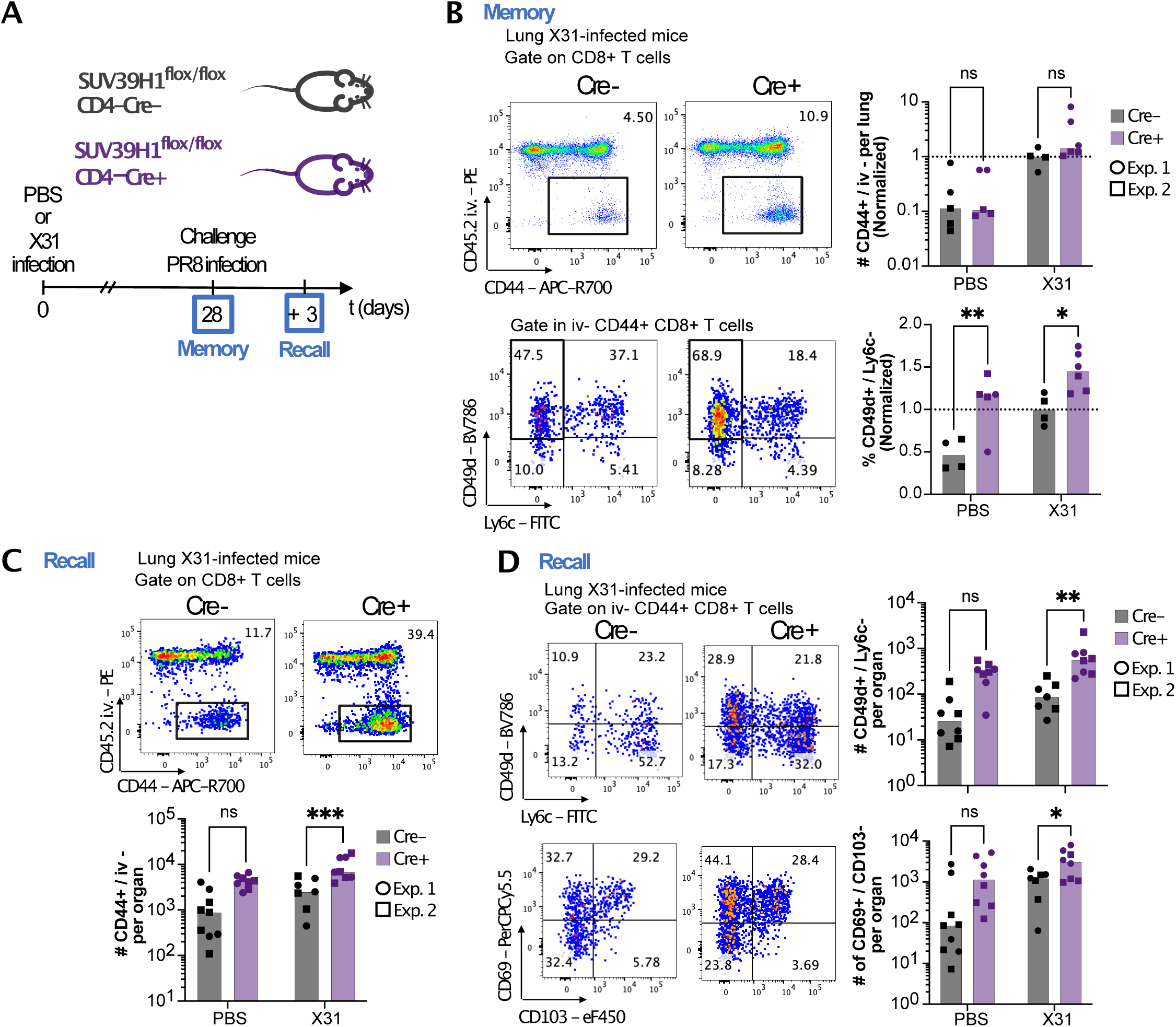
CD8^+^ T_RM_ differentiation upon flu infection is enhanced in the absence of SUV39H1. **A.** Scheme of the experiment. **B.** Representative dot plots and quantification of the normalized numbers of iv-CD44^+^ CD8^+^ T cells and the normalized percentages of CD49d^+^ Ly6c^-^ cells among iv-CD44^+^ CD8^+^ T cells in lungs 28 days post X31 infection (**Memory**). Values were normalized to the mean of X31-infected WT group for each experiment and then pooled. **C.** Representative dot plots and quantification of the number of iv-CD44^+^ CD8^+^ T cells in lungs 3 days post PR8 infection (**Recall**). **D.** Representative dot plots and quantification of the number of CD49d^+^ Ly6c^-^ and CD69^+^ CD103^-^ iv-CD44^+^ CD8^+^ T cells in lungs 3 days post PR8 infection (**Recall**). * p<0.05 ; ** p<0.01 ; *** p< 0.001 ; ns: not significant by ANOVA test with Šídák’s multiple comparisons test.

### CITE-seq analysis of lung-homing CD8^+^ T cells

To further characterize SUV39H1-KO T cells homing to tissues, we used CITE-seq analysis of sorted cells from lungs of RAG-KO mice that had received WT and SUV39H1-KO OT-I cells (as in Fig 2). To obtain sufficient cells for the analysis of all populations, we mixed equal numbers of cells from WT and KO genotypes (expressing distinct congenic markers CD45.1 and 2), coming from the iv+ or iv-fractions (Supp Fig 3 A). We used hashtag-coupled antibody staining to deconvolute the 4 populations (WT iv+, WT iv-, KO iv+, KO iv-), and we also stained all cells with hashtag-coupled antibodies for CD62L, CD44, CD49d, Ly6c, CD69 and CD103 to evaluate protein expression of these markers. We then performed scRNAseq using the 10x Genomics Chromium platform. In total, we obtained 4807 cells after removing non-T cell contaminants, low-quality cells, doublets, and cells with no hashtag identification. Unsupervised clustering partitioned cells into 9 clusters based on their transcriptome (Fig. 4 A). We used differential gene expression analyses and analyses of canonical marker genes (Fig 4 B) and gene signatures (Supp. Fig 3 B) to assign identities to the different clusters. Clusters were named after their class and highly expressed genes. As shown in Fig 4 A, we identified one cluster of cycling cells (Cycling Mki67), four clusters of central memory cells (T_CM_ CCR7 a-c and T_CM_ Jun), three clusters of effector cells (T_Eff_ Lgals3, CTL GzmA and T_Eff_ CD49d), and one cluster of tissue-resident memory cells (T_RM_ CD69, Fig 4 A). T_CM_ cells were separated from T_Eff_ and T_TAM_ cells on the Uniform Manifold Approximation and Projection (UMAP) dimensional reduction plot (Fig 4 A).

**Figure 4:**
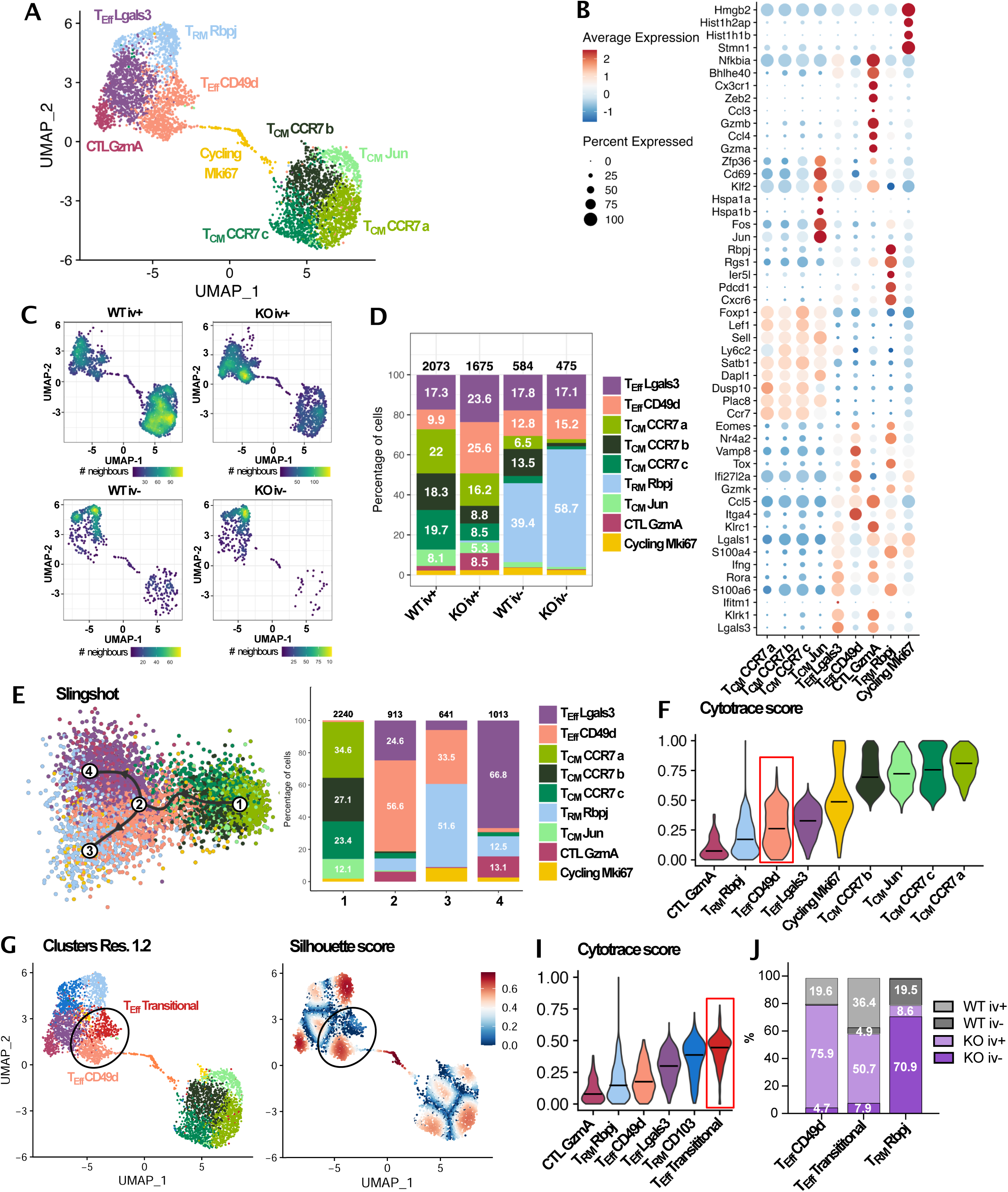
SUV39H1 regulates the developmental pathway of T_RM_ cells. OTI cells from lungs of 6 pooled mice from the experiment described in figure 2 A were FACS sorted as shown in Figure 2 C (WT iv+, KO iv+, WT iv-, KO iv-), stained with hashtag-labeled antibodies and pooled before performing 10X single cell CITEseq libraries. **A.** UMAP representation of the single cell RNAseq data showing clusters at resolution 0.8. **B.** Dot plot showing the top 5 differentially expressed genes in each cluster. **C.** UMAP representation of the single cell RNAseq data showing density scores of the 4 sorted samples. **D.** Distribution of clusters in each sample. Numbers within the bar plots represent the percentage of indicated clusters in each sample. Numbers above bar plots indicate the absolute number of cells for each sample. **E.** UMAP representation of Slingshot trajectory analysis on RNAseq data showing 4 milestones (**left**) and distribution of each cluster in the 4 milestones (**right**). Numbers within the bar plots represent the percentage of indicated clusters in each sample. Numbers above bar plots indicate the absolute number of cells for each sample. **F.** Violin plots showing Cytotrace stemness score distribution for each cluster. **G.** UMAP representation of the single cell RNAseq data showing clusters at resolution 1.2. **H.** UMAP representation of the single cell RNAseq data showing Slhouette scores at resolution 1.2. **I.** Violin plots showing Cytotrace stemness score distribution for the indicated clusters at resolution 1.2. **J.** Computing of original proportions of samples in each cluster (See Methods). Numbers represent proportion of each sample within each cluster. Each dot represents a cell.

The top 5 upregulated genes in T_CM_ clusters (T_CM_ CCR7 a-c and T_CM_ Jun) include Foxp1, Sell, CCR7 and Ly6c2 -encoding Ly6c, except for T_CM_ CCR7 a-(Fig 4 B). Cluster T_CM_ Jun also expresses Fos and CD69, probably reflecting recent activation. This cluster also upregulated Klf2, a transcription factor expressed in recirculating cells^35^ (Fig 4 B). The cytotoxic CTL GzmA cluster expresses markers of cytotoxic effectors, like granzyme A and B (Gzma, Gzmb), CX3CR1 and interferon gamma (Ifng). Also, effector cluster T_Eff_ Lgals3 expresses effector (Ifng) and terminal effector markers such as Klrc1 and Klrk1, encoding NKG2D (Fig 4 B). In addition, we found Itga4 (encoding CD49d) among the top 5 upregulated genes in effector cluster T_Eff_ CD49d. This cluster also expresses high levels of effector and memory markers such as Ccl5, Eomes and Tox as well as Gzmk and Nr4a2. Among these genes, some are also implicated in T_RM_ differentiation, such as Nr4a2^11^, or T_RM_ maintenance, such as Eomes^36^.

Additionally, some of these genes are also upregulated upon TCR-dependent cell activation, like Tox and Eomes^37,38^ (Fig 4 B). Among the top 5 upregulated genes in T_RM_ CD69 cells are Rbpj^32^, Cxcr6^12^ and Pdcd1 (encoding PD1^15^), all markers of bona fide T_RM_ cells, and RGS1, a G protein regulator that is involved in inhibition of T cell trafficking to lymph nodes^39^ (Fig 4 B). Additionally, cluster T_RM_ CD69 expresses high levels of T_RM_ signatures from the literature (Supp Fig 3 B). Other typic markers of T_RM_ cells like Hobbit (Zfp683), Blimp-1 (Prdm1) or CD103 (Itgae) were not detected at the RNA level in our dataset. To further evaluate the identity of this cluster, we analyzed transcription factors (TF) activity using Single-Cell rEgulatory Network Inference and Clustering (SCENIC^40^, Supp Fig 3 C). Among the 10 highest fold-changes in TF activity (regulons) for each cluster, we identified several active regulons in T_RM_ CD69 cells that are related to T_RM_ biology. Among these regulons is Srebp2, recently involved in the metabolic adaptation of T_RM_ cells^41^; the AP-1 family members Fosb, Fos, Jund and Jun, highly expressed in T_RM_ cells^15,32,42^; and Prmd1^12^ -encoding for BLIMP1-Runx3^8^, Tbx21^43^ -encoding for Tbet-, and Irf4^44^ all having roles in T_RM_ formation and maintenance.

Regarding the expression levels of our protein markers, T_RM_ CD69, T_Eff_ CD49d, as well as T_Eff_ Lgals3, and Cycling Mki67 clusters, all express high levels of CD49d and low levels of Ly6c proteins, as detected by hashtag-coupled antibodies (Supp Fig 3 D and E). In addition, even though Cd69, encoding CD69, was expressed at similar RNA levels in all clusters, except for T_CM_ Jun cluster that expressed higher levels (Supp Fig 3 F), it was highly expressed at the protein level only in cluster T_RM_ CD69 (Supp Fig 3 E), consistent with previous reports of discrepancy of RNA and protein expression for this gene in the hematopoietic compartment^45^. Finally, as expected, cluster T_RM_ CD69 is specific of lung extravascular compartments (iv-)(Fig 3 C and D). Consistent with previous results, both T_TAM_ CD69 and T_Eff_ CD49d clusters were among the most overrepresented in KO cells (together with the minor cytotoxic cluster CTL GzmA), compared to WT cells (Fig 4 C, D and Supp Fig 3 G). Altogether, these results show that SUV39H1 deficiency leads to accumulation of CD8^+^ T cells in lung tissue, suggesting that epigenetic silencing by SUV39H1 restrains tissue homing of CD8+ T cells and the differentiation of T_RM_ cells in a cell-intrinsic manner.

### SUV39H1 regulates the developmental pathway of T_RM_ cells

To investigate a possible relationship between T_Eff_ cells expressing CD49d and T_RM_ cells accumulating among SUV39H1-defficient cells, we performed trajectory analysis using 2 different algorithms, Slingshot^46^ and Scvelo^47^. Both analyses suggested a differentiation trajectory starting with T_CM_ CCR7 clusters, which differentiate into T_Eff_ CD49d. T_Eff_ CD49d represent a branching point to the two final differentiation branches that would then drive to either T_Eff_ Lgasl3 and CTL GzmA or T_RM_ CD69 (Fig 4 E, Supp Fig 4 A). The majority of the top 50 genes varying the most along the trajectory ordered by Slingshot pseudo-time, are shared between T_RM_ CD69 and the three effector clusters (T_Eff_ Lgals3, T_Eff_ CD49d and CTL GzmA; Supp Fig 4 B). Some of these genes (CD3g, Itga4, GzmK and Tox) are shared by T_Eff_ CD49d and T_RM_ CD69, but not by the other effector clusters. Altogether, these results suggest that T_Eff_ CD49d cells are enriched in precursor cells that could give rise to T_RM_ CD69 cells. This hypothesis is consistent with most cycling cells being related to T_Eff_ CD49d cluster, as suggested by label transfer of cycling cells into other clusters (using Seurat label transfer method, Supp Fig 4 C).

To evaluate if the T_Eff_ CD49d cluster is enriched in precursors cells, we used the Cytotrace score, a proxy for stemness as a function of gene expression diversity per cell^48^. As shown in Fig 4 F, the Cytotrace score for cluster T_Eff_ CD49d showed bimodal distribution, suggesting the presence of a high stemness subcluster among T_Eff_ CD49d cells. To further explore this possibility, we computed the Silhouette score for each cell, a parameter that indicates how well assigned to its cluster each cell is. This score can be used to estimate if a cluster represents a defined stable cluster (high Silhouette score), or a transitional state between two stable clusters (low Silhouette score). At 0.8 resolution, all clusters have high Silhouette scores (Supp Fig 4 D). Nevertheless, at higher resolution (1.2), T_Eff_ CD49d, resolved into 2 clusters, one of which has lower Silhouette score than all other clusters (Fig 4 G and H), suggesting that these cells are transitional (i.e. in transition between two stable states). This transitional cluster share upregulated genes with T_Eff_ CD49d cells, including Itga4, Eomes and Nr4a2 (Supp Fig 4 E). Some of the upregulated genes in the T_Eff_ transitional cluster, including Eomes and Slamf6, are also upregulated in T_RM_ precursors^11^.

Slingshot analysis at this higher resolution indicates that T_Eff_ transitional cluster differentiates into T_RM_ clusters (T_RM_ CD69 or T_RM_ CD103) and circulatory effector clusters (T_Eff_ Lgals3 and CTL GzmA, Supp Fig 4 F). Likewise, Cytotrace analysis at 1.2 resolution showed that the T_Eff_ transitional cluster has higher stemness score than all other effector and T_RM_-like clusters (Fig 4 I). Altogether these results indicate that the T_Eff_ CD49d cluster includes transitional cells with features of T_RM_ cell precursors. Even though cluster T_Eff_ transitional was not overrepresented in SUV39H1-deficient samples (Supp Fig 4 G), computation of the original proportions of cells in lungs before sorting (see Methods) showed that the majority of T_Eff_ transitional cells in lungs were deficient for SUV39H1 (Fig 4 J). Altogether our results suggest that CD49d-expressing T_RM_ precursors accumulate in the absence of SUV39H1 expression.

### T_RM_ precursors differentiation is increased in the absence of SUV39H1

To investigate if cells that accumulate in tissues in the absence of SUV39H1 have increased T_RM_ precursor potential, we transferred WT and SUV39H1-defficeint naïve OT-I cells in a 1:1 ratio into B6 mice and then infected the mice with the flu strain X31 expressing the SIINFEKL peptide (X31-OVA, Fig 5 A). 30 days after infection, CD69^+^ CD103^-^ T cells accumulated among SUV39H1-KO OT-I cells in lung tissue (Fig 5 B and Supp Fig 5 A), reminiscent of what we have seen ex-vivo at steady state (Fig 1) and after flu infection (Supp Fig 3). We also found an increase in CD69^+^ CD103^+^ numbers, but not proportions, among SUV39H1-deficient OT-I cells compared to WT control cells (Fig 5 B). SUV39H1-KO cells expressing CD49d accumulated in the lungs during the effector phase of the infection (Fig 5 C and Supp Fig 5 B). Also, cells expressing CD49d and low levels of Ly6c were enriched in CD69^+^ KLRG1^-^ and CD49a^+^ CX3CR1^-^ cells, both expressed in T_RM_ precursors^8,9^ (Fig 5 C and Supp Fig 5 C). We also observed an increase in CD49a+ CX3CR1-cells among liver-homing CD8^+^ T cells at steady state (Supp Fig 5 D). Altogether these results suggest that SUV39H1-defficiency leads to accumulation of T_RM_ precursors both at steady state and during the effector phase of flu infection.

**Figure 5:**
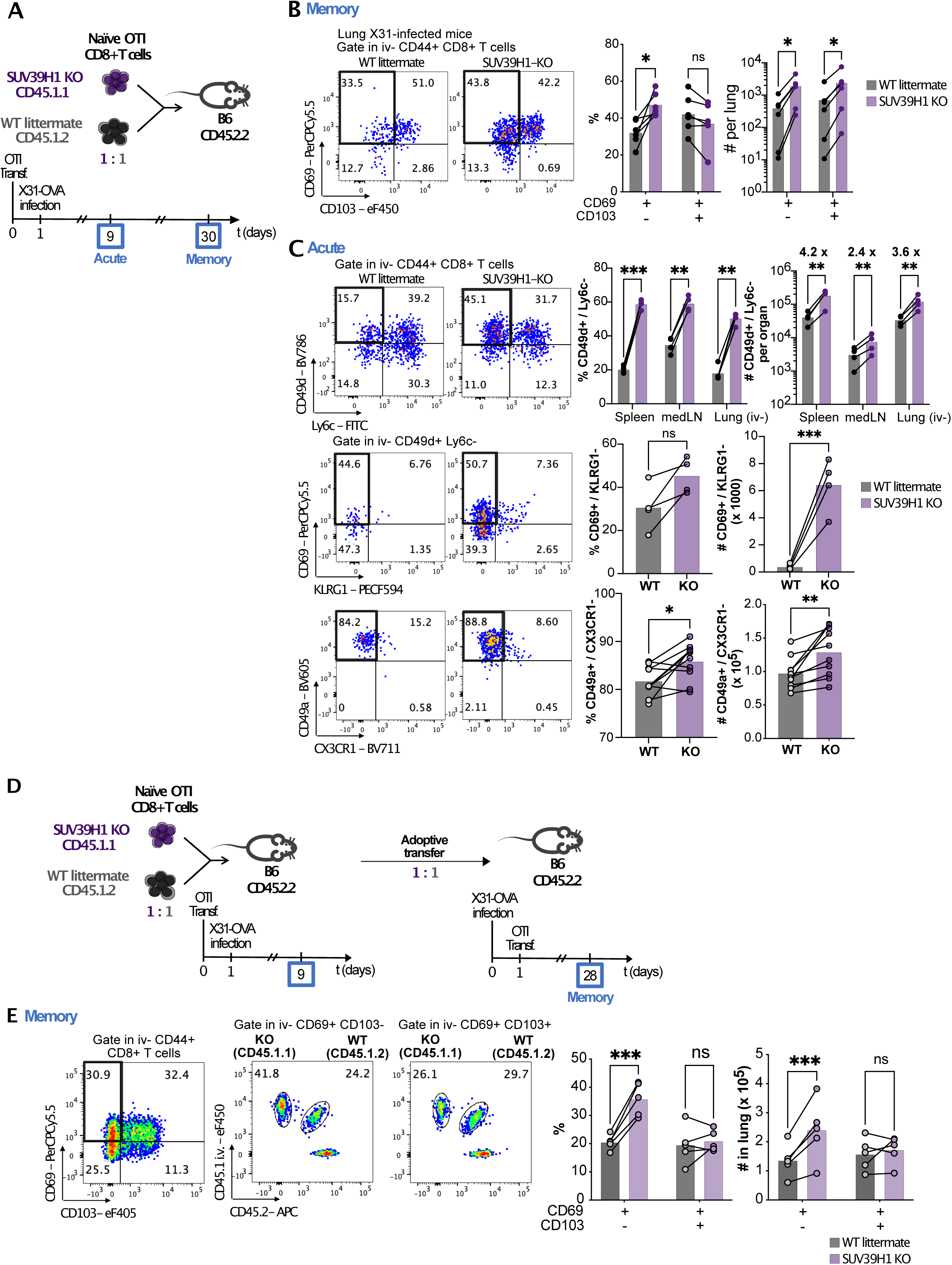
T_RM_ precursors differentiation is increased in the absence of SUV39H1. **A.** Scheme of the experiment. **B.** Representative dot plots and quantification of the expression of CD69 and CD103 in SUV39H1-deficient and WT OTI cells 30 days post X31-OVA infection in lungs (**Memory**). **C.** Representative dot plots from lungs and quantification of the percentage of CD49d^+^ Ly6c^-^ cells among SUV39H1-deficient and WT OTI cells in the indicated organs and the percentage of CD69^+^ KLRG1^-^ or CD49a^+^ CX3CR1^-^ cells among CD49d^+^ Ly6c^-^ iv-CD44^+^ OTI cells in lungs 9 days post X31-OVA infection (**Acute**). **D.** Scheme of the experiment. **E.** Gating strategy, representative dot plots and quantification of the percentage and numbers of WT and SUV39H1 KO OTI cells among CD69^+^ CD103^-^ and CD69^+^ CD103^+^ iv-CD44^+^ OTI cells in lungs 28 days post X31-OVA infection (**Memory**) and 27 days post adoptive transfer. p<0.05 ; ** p<0.01 ; *** p< 0.001 ; **** p<0.0001 ; ns: not significant by multiple ratio paired t test (B), (C) top and (E) and ratio pared t test (C) bottom. **B, C and E**: Representative data from 2 independent experiments.

To formally test if SUV39H1 KO enhances T_RM_ precursor function, WT and KO OTI cells from the intravascular (iv-) compartment of the lung were FACS-sorted 9 days after X31-OVA infection and adoptively transferred intravenously at 1:1 ratio into recipient mice previously infected with X31-OVA (Fig. 5 D and Supp Fig 5 E). After 28 days, mice had higher proportions of SUV39H1-deficient OT-I cells, compared to WT cells, among CD69^+^ CD103^-^ CD8^+^ T cells in the iv-compartment of the lung (Fig 5 E). These findings show that SUV39H1-deficient CD8^+^ T cells homing to the lungs during the acute phase of infection are enriched in CD69-expressing T_RM_ precursors compared to WT cells.

### SUV39H1-deficient lung-homing CD8^+^ T cells delay lung tumor metastasis growth

The results presented so far show that SUV39H1 deficiency increases T_RM_ precursor frequency and tissue colonization in lungs, both, after Flu infection (Fig 3) and during spaced-induced expansion in RAG KO mice (Fig 2). To evaluate the protective potential of the tissue-homing CD8^+^ T cells formed in the absence of SUV39H1, we transferred naïve OT-I CD8^+^ T cells from SUV39H1-KO mice or WT littermates into RAG-KO mice. Between 33 and 35 days later, we challenged the mice with an intravenous injection of B16-OVA cells expressing Luciferase (B16-OVA-Luc, Fig 6 A). The numbers of iv-CD8^+^ T cells in the lungs 33 days after transfer were similar in mice that received WT or SUV39H1-KO OT-I cells (Fig 6 B), while SUV39H1-KO iv-cells were enriched in CD49d^+^ Ly6c^-^ cells, reminiscent of what we found in co-transfer experiments. Mice adoptively transferred with either WT or SUV39H1-KO OT-1 cells showed delayed B16-OVA-Luc growth as compared to mice that did not receive any OTI cells (Fig 6 C). In mice that received SUV39H1-KO OT-I cells, however, tumor growth delay was significantly stronger compared to mice bearing WT OT-I cells (Fig 6 C and D).

**Figure 6.**
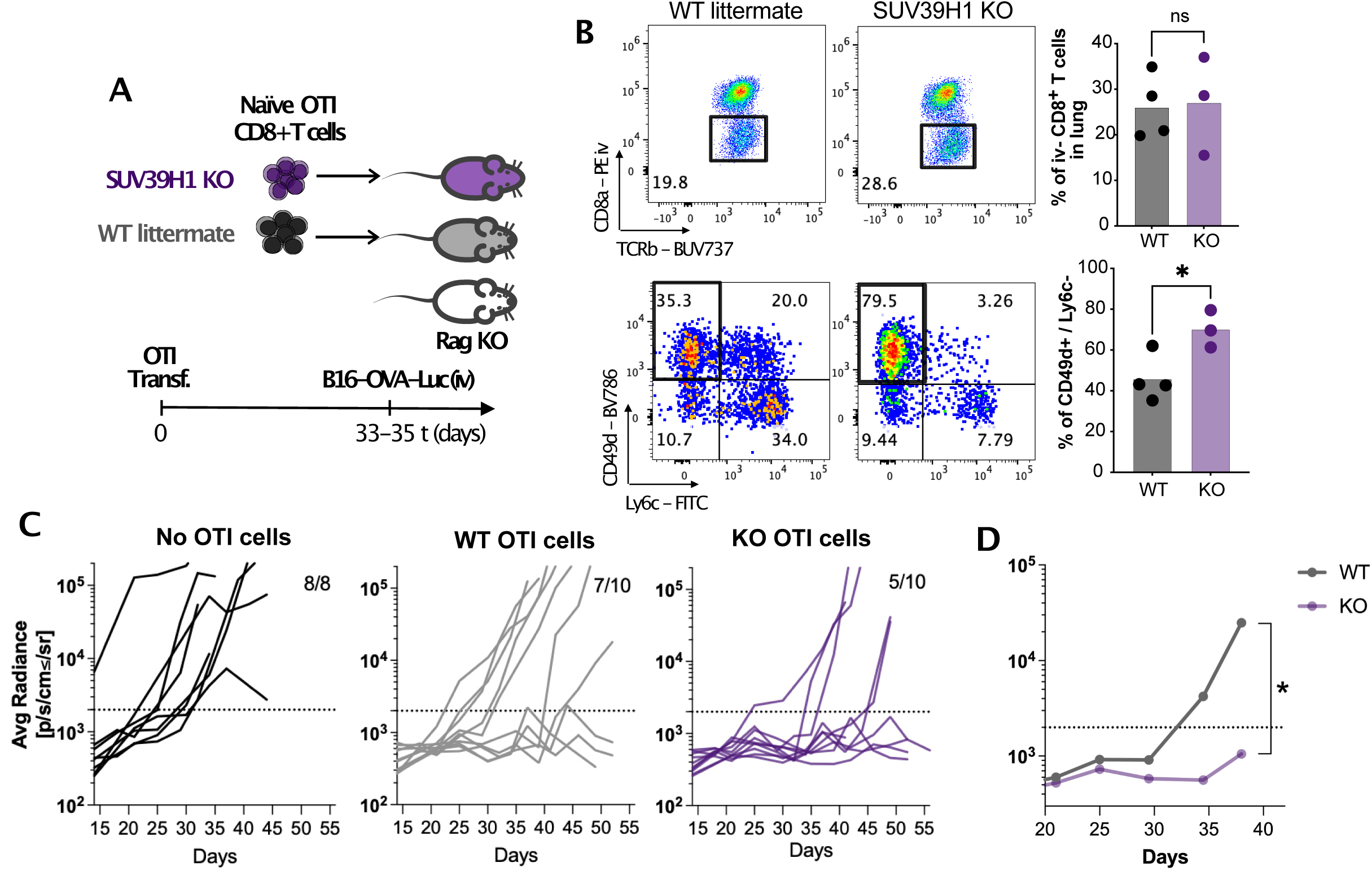
SUV39H1-deficient lung-homing CD8+ T cells delay lung tumor metastasis growth. **A.** Scheme of the experiment. **B.** Representative dos plots of iv staining (**top**) and CD49d and Ly6c expression in SUV39H1 KO and WT iv-OTI cells (**bottom**) in the lungs of mice at 33 days post transfer. **C.** Individual tumor growth curves in each experimental group. Inset ratios show the number of mice that developed tumors at the end of the experiment over the total number of mice in each experimental group (cut off average radiance = 2×10^3^ p/s/cm^2^/sr). Pooled of 2 independent experiments. **D.** Median tumor growth curves of SUV39H1 KO and WT OTI recipient mice in **C**. Each dot represents a mouse. * p<0.05; ns: not significant by unpaired t test (**B**) and mixed-effects model (**D**).

These results suggest that, even though the proportions of CD8^+^ T cells in the lung tissue were similar in WT and SUV39H1 KO mice, the increased proportions of CD8^+^ CD49d^+^ T cells formed in the absence of SUV39H1 are more efficient delaying tumor growth, compared to WT CD8^+^ T cells.

## Discussion

The molecular mechanisms that lead to the establishment and maintenance of effective immunological memory are still incompletely understood. The memory T cell compartment is indeed particularly complex, with multiple types of memory cells in lymphoid and non-lymphoid tissues. Most T cell memory models propose that memory precursor T cells differentiate early during T primary immune responses in lymph nodes and survive the contraction of effector populations that follows pathogen clearance^8,49–51^. These long-lived memory precursors either stay in lymphoid tissues and become T_CM_ and T_EM_ (which maintain their circulatory potential), or colonize tissues, where they further differentiate into T_RM_ (that do not recirculate).

We show here that like T_CM_ in lymphoid tissues, CD8^+^ T cell numbers and proportions are increased in non-lymphoid tissues in absence of SUV39H1 expression. Increased presence of CD8^+^ T cells occurs in different tissues, including lungs, liver, and small gut. It occurs at steady state, and after spaced-induced “homeostatic” expansion or Flu infection. Then, what is the cellular mechanism underlying these differences: increased tissue colonization, T cell lifespan and /or tissue retention? The effect of the SUV39H1-defect was particularly strong in a model of competition between WT and KO T cells, in which SUV39H1 KO CD8^+^ non-lymphoid tissue-homing cells surpassed WT T_RM_ by 10-folds in small gut or liver colonization.

In addition, scRNAseq analyses showed increased proportions of a population of T_RM_ precursors in SUV39H1 KO cells homing to lungs compared to WT cells. Adoptive transfer of purified SUV39H1-defficient lung homing effectors to Flu infected mice shows increased T_RM_ colonization, indicating increased T_RM_ precursor capacity in absence of SUV39H1 expression. Altogether, these results suggest that CD8^+^ T cells, including T_RM_ precursors, colonize non-lymphoid tissues more efficiently in absence of SUV39H1 expression. Consistent with this interpretation, we have not observed evidence that SUV39H1 KO affects T cell survival or expansion in tissues (no differences in apoptosis signatures or in dividing T cell populations in the single cell analyses). Increased tissue colonization is also consistent with the observed increased expression of CD49d in SUV39H1 KO T cells, necessary for the trans-migration of T cells from blood to tissues^52^.

At the molecular level, the mechanism involved is less clear. Our previous studies showed that critical memory differentiation genes, including IL-7R, are silenced by H3K9me3 methylation during effector differentiation^19^. H3K9me3 levels at these loci are reduced in SUV39H1 KO T cells, indicating a direct role for SUV39H1 in the silencing of these memory-related genes. These results suggest that H3K9 trimethylation by SUV39H1 plays a critical role in the control of the balance between effector and memory differentiation. The results presented here could derive from a similar mechanism for SUV39H1 in the control of T_RM_ differentiation: some of the loci that are silenced by SUV39H1 during the differentiation of short-lived effectors, like IL-7R, could promote T_RM_ differentiation in absence of SUV39H1 expression. Therefore, our results suggest that the epigenetic pathways that control T_CM_ and T_RM_ share some fundamental control mechanisms.

Can we then conclude that T_CM_ and T_RM_ have common precursors? Our results have not addressed this possibility directly, but adoptive transfer of SUV39H1-deficient cells in the peak of the infection have both higher potentials to become T_RM_ (Fig 5 D) and T_CM_^19^. Studies from the Mackay lab and others show that T_RM_ arise from a pool of memory precursors effector cells that resists contraction after resolution of infection^10,51^. Even though the regulation of lineage choice between circulating and resident memory cells is not completely understood, some T_RM_-associated molecules like CD69 are expressed very early in effector cells located into tissues, suggesting that T_RM_ and circulating memory cells linages separate early on^8,51^. SUV39H1 ablation, we show here, increases both tissue accumulation and the differentiation of T_RM_ precursors among tissue homing-effector cells, causing T_RM_ cells accumulation in tissues. These T_RM_ precursor cells express high levels of the integrin CD49d and other molecules found in T_RM_ precursors by others, including CD49a, Slamf6, EOMES, CD69 and IL7R^9–11^. The results presented here suggest that T_RM_ and T_CM_ precursors share epigenetic mechanisms of differentiation. This could be related to a common origin of T_CM_ and T_RM_ precursors. Alternatively, stemness gene repression in effector cells by SUV39H1 could counteract both T_CM_ and T_RM_ precursors independently.

T_RM_ populations in tissues are heterogenous^3,53^. CD103^+^ CD8^+^ T_RM_ cells locate to epithelial structures, while CD103^-^ T_RM_ cells home to the stroma^54^. Also, recent work shows that CD103-T_RM_ cells can replenish the CD103+ T_RM_ pool upon re-infection^55–57^. The T_RM_ accumulation observed in absence of SUV39H1 expression mostly concerns CD103^-^ T_RM_, suggesting that SUV39H1 KO can promote the differentiation of CD69^+^CD103^-^ or reduce the transition to CD103^+^ T_RM_ cells. In any case, over representation of CD49d^+^ Ly6c^-^ T_RM_ in SUV39H1-KO is consistent with increased tissue homing (due to increased CD49d expression) and reduced lymphoid homing (due to reduced Ly6c expression).

Several studies focusing on epigenetic changes during T_RM_ differentiation have described both common and tissue specific regulatory networks and key transcription factors directing the generation and maintenance of the T_RM_ linage^15,58,59^. However, the role of specific epigenetic regulators and chromatin modifiers in T_RM_ formation is largely unknown. McDonald et al, found that the canonical BAF complex, crucial for nucleosome reorganization during novel enhancer establishment, was necessary for late-stage differentiation of T_RM_ cells but was dispensable for circulatory memory formation^60^. The role of SUV39H1 in T_RM_ differentiation, seems to be the opposite to BAF. It is possible that SUV39H1 is necessary to silence specific enhancers during effector differentiation, leading to enhanced T_CM_ and T_RM_ differentiation in absence of SUV39H1.

Previous work from our lab also shows that SUV39H1-deficiency in CAR-T cells improve anti-tumor efficacy^28^. This effect was linked to less exhaustion and higher memory persistence in the absence of SUV39H1. On the contrary, the capacity of memory SUV39H1-deficient cells to prevent bloodborne *Listeria monocytogenes* re-infection was reduced compared to WT cells^19^. Here we show that CD8^+^ T_RM_ formed in the absence of SUV39H1 show a relatively higher efficacy to control pulmonary flu-reinfection. Also, SUV39H1-deficient CD8+ T cells homing to lungs were able to longer delay tumor metastases growth compared to WT cells. Even though we cannot exclude an effect of circulatory cells in the higher protection against virus and tumors, the results presented here suggest that the increase in tissue homing and the subsequent increased T_RM_ cell differentiation in absence of SUV39H1 improves the efficacy of memory immune responses in tissues, both against tumors and infections. Inhibition of expression or function of SUV39H1 in T cells should help promote efficient immune responses in the context of both adoptive and non-adoptive immunotherapies.

## Methods

### Mice, virus strains and cell lines

C57BL/6J male mice were obtained from Charles River and used at 6-10 weeks of age. Suv39h1^tm1Jnw^C57BL/6J (Suv39h1-KO) mice kindly provided by T. Jenuwein and previously described^61^, were backcrossed with C57Bl6/J mice for at least nine generations. C57BL/6J-Tg(TcraTcrb)1100Mjb/J mice (OT-I) mice and B6.SJL-Ptprc^a^ Pepc^b^/BoyJ (CD45.1) mice were purchased from Jackson lab. OT-I mice and CD45.1 mice were crossed to obtain OT-I/CD45.1/CD45.2 mice. B6-Suv39h1^tm1Ciphe^ (Suv39h1^Flox/Flox^) originally produced at the Centre d’Immunologie de Marseille (CIPHE, B. Malissen) and previously described^29^ were crossed with B6;D2-Tg(Cd4-cre)1Cwi/CwiCnrm (CD4-Cre) mice obtained from the European Mouse Mutant Archive to obtain Suv39h1^Flox/Flox^ CD4-Cre mice. CD3 KO mice were obtained from B. Malissen^62^, Rag-KO mice were obtained from Charles River. Suv39h1Flox*CD4-Cre+/− and control Suv39h1-Flox*CD4-Cre−/− mice were used for the experiments. Male mice, of either Suv39h1-KO, OT-1/CD45.1 mouse strains were used aged 6–12 weeks. Male and female mice of Suv39h1^Flox/Flox^ CD4-Cre mouse strain were used aged 6–12 weeks unless otherwise stated.

All mice were housed in Curie Institute SPF animal facilities. Live animal experiments were performed in accordance with the guidelines of the Ministere de l’enseignement superieur, de la recherche et de l’innovation (Project authorization: APAFIS #34372-2021121522311823 v1). Influenza A/HKX31 (H3N2) and A/PR8 (H1N1) virus were purchased from Charles River. Influenza A/HKX31 (H3N2) and A/PR8 (H1N1) virus expressing the OVA peptide SIINFEKL, (X31-OVA and PR8-OVA, respectively) were kindly provided by Dr. Stephen Turner and Dr. Richard Webby^63^. B16F10-expressing OVA (B16F10-OVA) stably expressing Luciferase (B16F10-OVA-Luc) were produced by transducing B16F10-OVA cells with lentiviral particles produced with the modified pTRIP-BFP-2a-RedLuc-pEF1-YP plasmid (kindly provided by Dr. Frank Perez), as previously described^63^.

### Intravascular staining and tissue preparation

Tissue homing and circulating cells were distinguished by intravascular staining as previously described^30^. Briefly, 15 ug of CD45.1-PE, CD45.1-APC, CD45.2-PE, CD45.2-APC, CD8α-PE or CD8α-APC in 100 ul of PBS were injected intravenously in the retroorbital sinus between 3’ and 5’ before killing the mice.

To isolate lymphocytes from the lungs, lungs were digested using GentleMACS® tissue dissociator (Milteny) in CO2 independent medium (Gibco 18045-054) with DNAse I 0,1mg/ml (Roche 11-284-932-001) and Liberase® 0,1 mg/ml (Roche 05-401-020-001).

For isolation of lymphocytes from the small intestinal epithelium (IELs), small gut was dissected from the caecum to the stomach, Peyer’s patches were removed, and the tissue was cut longitudinally and washed of luminal contents. The tissue was then cut into 0.5-cm pieces that were incubated while shaking in Hanks’ balanced salt solution (HBSS) without Ca2+ and Mg2+ (Gibco 14185-052) with EDTA 5 mM (Invitrogen 15575-038), FBS (Eurobio) 10% and DTT 1 mM (Euromedex 0006-b) at 37°C 3 times for 20’ and 1 time in HBSS without Ca2+ and Mg2+ for 20’. Cells within the supernatant were collected.

To isolate lymphocyte from the small gut Lamina Propria (LP) remaining gut pieces were incubated at 37 C with agitation in 2.5 ml of CO2 independent medium (Gibco 18045-054) with 0.17 U/ml of Liberase® (Roche 05-401-020-001), 1 mg/ml of collagenase (Sigma-Aldrich 11088866001) and 125 ug/ml of DNAse I (Roche 11284932001) for 30’ and then mechanically disaggregated in gentleMACS® tissue dissociator.

To isolate lymphocytes from skin, skin from the back of the mice was shaved and cleaned with EtOH. Then subcutaneous fat tissue was separated from the skin with the help of scalpels. Cleaned skin was then cut in small pieces with scissors and digested in 6 ml of CO2 independent medium (Gibco 18045-054) containing Hyaluronidase type IS 500 µg/ml (Sigma-Aldrich H3506), Liberase® 100 µg/ml (Roche 05-401-020-001), DNAse I 250 µg/ml (Roche 11284932001), 2% FBS (Eurobio) and 1% Hepes (Gibco) for 1h at 37 C with agitation. Then, tissue was mechanically disaggregated by using gentleMACS® tissue dissociator.

Liver was scratched using a 100 µM cell strainer (Falcon 352360) and a syringe plunge in PBS (Eurobio CS1PBS01-01) supplemented with EDTA 2 mM (Invitrogen 15575-038) and BSA 0.5% (Sigma-Aldrich A7906) -FACS buffer-.

Lymphocytes from digested lung, IELs, LP, skin and liver samples were enriched by 40%/80% Percoll® (Sigma-Aldrich) gradient^64^.

Mediastinal, inguinal, and mesenteric lymph nodes and spleen were scratched using a 40 µM cell strainer (Falcon 352340) and a syringe plunge in CO2 independent medium (Gibco 18045-054) supplemented with 10% FBS (Eurobio).

### Cell staining and flow cytometry analysis

Cell suspensions were prepared in PBS and stained with Near IR (Fisher Scientific L34975) or Acqua LIVE/DEAD™ Fixable Cell Stain Kit (Fisher Scientific L34957) according to manufacturer’s instructions to exclude dead cells and then labelled in FACS buffer with the following antibodies: V500 Rat anti-Mouse CD4 (RM4-5, BD), eF450 CD5 Monoclonal Antibody (53-7.3, eBiosciences), FITC anti-CD8α (53-6.7, BD), PerCP-Cy5.5 CD8 α (53-6.7, BD), PE CD8 α Monoclonal Antibody (53-6.7, Ficher Scientific), BUV395 Rat Anti-Mouse CD8 β (H35-17.2, BD), APC anti-mouse CD25 Antibody (PC61, Biolegend), PerCP-Cy5.5 Rat Anti-Mouse CD44 (IM7, BD), APC Rat Anti-Mouse CD44 (IM7, BD), APC-R700 Rat Anti-Mouse CD44 (IM7, BD), eFluor 450 CD45.1 Monoclonal Antibody (A20, Fisher Scientific), PE Mouse Anti-Mouse CD45.1 (A20, BD), APC-Cy7 CD45.1 (A20, BD), PE Mouse Anti-Mouse CD45.2 (104, BD), APC Mouse anti-Mouse CD45.2 (104, BD), BV605 Hamster Anti-Rat/Mouse CD49a (Ha31/8, BD), BV786 Rat Anti-Mouse CD49d (R1-2, BD), PE/Cyanine7 anti-mouse CD49d Antibody (R1-2, Biolegend), PE-Cy7 Rat Anti-Mouse CD62L (MEL-14, BD), APC Rat Anti-Mouse CD62L (MEL-14, BD), APC/Cyanine7 anti-mouse CD62L Antibody (MEL-14, Biolegend), PerCP-Cy5.5 Hamster Anti-Mouse CD69 (H1.2F3, BD), PE-Cy7 Hamster Anti-Mouse CD69 (H1.2F3, BD), Brilliant Violet 605™ anti-mouse CD73 Antibody (TY/11.8, Biolegend), eFluor 450 CD103 Monoclonal Antibody (2E7, Fisher Scientific), eF660 CD127 (A7R34, eBiosciences), Brilliant Violet 711 anti-mouse CX3CR1 (SA011F11, Biolegend), BV711 CXCR3 (CXCR3-173, BD), Brilliant Violet 650™ anti-mouse Ly-6C Antibody (HK1.4, Biolegend), FITC anti-mouse Ly-6C (HK1.4, Biolegend), APC Ly-6C Monoclonal Antibody (HK1.4, eBiosciences), PE-CF594 Hamster Anti-Mouse KLRG1 (2F1, BD), BUV737 Hamster Anti-Mouse TCR β Chain (H57-597, BD), Alexa-Fluor®488 Hamster Anti-Mouse TCR β Chain (H57-597, BD), PE-Cy™7 Hamster Anti-Mouse TCR β Chain (H57-597, BD), PE-CF594 TCR β Chain (H57-597, BD), eFluor 450 TRCR V alpha 2 (B20.1, Invitrogen), PerCP/Cy5.5 anti-mouse TCR Vα2 (B20.1, Biolegend).

For some experiments, cells were fixed in 2% PFA (Electron Microscopy Sciences) during 20’ at room temperature. Cell suspensions were analyzed in a BD FacsVerse or a BD LSRFortessa cytometer. Data was analyzed by Flowjo 10 software version: 10.8.0.

### Adoptive cell transfer

Secondary lymphoid organs (spleen, mesenteric, axillary, inguinal, brachial, cervical, and lumbar lymph nodes) from SUV39H1-KO or WT OT-I male mice were collected, pooled, and homogenized into single cell suspensions. Naïve (CD44-) CD8^+^ T cells were enriched using the naïve CD8_+_ T-cell isolation kit (Miltenyi 130-096-543) according to manufacturer’s instructions. 1 × 10^6^ or 5×10^4^ cells of a 1:1 mix of Suv39H1 KO and WT naïve CD8^+^ T cells were adoptively transferred into Rag-KO or C57BL/6J male mice respectively.

Adoptive transfer of T_RM_ precursors was made from lung tissue. Mice were injected with PE CD8 α Monoclonal Antibody (53-6.7, Ficher Scientific) as described before euthanasia. Lungs were digested as previously described and pooled enriched lymphocytes were stained with Acqua LIVE/DEAD™ Fixable Cell Stain Kit (Fisher Scientific L34957), eFluor 450 CD45.1 (A20, Fisher Scientific), PerCP-Cy5.5 CD8 α (53-6.7, BD), FITC anti-mouse Ly-6C (HK1.4, Biolegend), PE-CF594 TCR β Chain (H57-597, BD), PE/Cyanine7 anti-mouse CD49d Antibody (R1-2, Biolegend), APC Mouse anti-Mouse CD45.2 (104, BD), APC-R700 Rat Anti-Mouse CD44 (IM7, BD), APC/Cyanine7 anti-mouse CD62L Antibody (MEL-14, Biolegend). WT or SUV39H1 KO Total iv-, CD49d+/Ly6c- or CD49d+/Ly6c+ populations were FACS sorted from pooled lungs. Between 2×10^4 and 4×10^4 cells of each population of a 1:1 mix of Suv39H1 KO and WT cells were adoptively transferred. Cells were transferred intravenously by retroorbital injection.

### Flu infection

Mice were infected by intranasal instillation with a sublethal dose of Influenza PR8 (5×10^5^ EID50), X31 (2×10^5^ EID50) or X31-OVA (2×10^5^ TCID50) in 20 or 40 ul of PBS in the left nostril. Control group mice were instilled intranasally with PBS. In some experiments, 28 days after the first infection, mice were infected with a lethal dose of PR8 (2×10^6^ -200-EID50).

All infections were performed on anesthetized mice with ketamine (100mg/kg, Merial) and xylazine (7,5mg/kg, Bayer) in PBS administered intraperitoneally. The humane endpoint was defined by a maximum of 30% weight loss and additional clinical signs of distress.

### Viral load by Q-PCR

Lungs were scratched using a 100 µM cell strainer (Falcon 352360) and a syringe plunge in FACS buffer and half of the lungs was centrifuged and lysed in RLT buffer (Qiagen 79216) supplemented with 1% BMeOH (Sigma-Aldrich M3148) and saved at -80 C until further processing. RNA was extracted using RNeasy Mini kit (Qiagen 74104). RT PCR was performed using Superscript III (Invitrogen 10432122) and random primers (Promega C118A) following manufacturer’s instructions. Specific TaqMan probe for PA protein was designed using the AlleleID software (Premier Biosoft) using default parameters. Custom TaqMan probe (CCAAGTCATGAAGGAGAGGGAATACCGCT) was ordered from Fisher Scientific, and primers (Forward Primer: CGGTCCAAATTCCTGCTGA; Reverse primer: CATTGGGTTCCTTCCATCCA) were from Eurogentec. Concentrated TaqMan assays (20×) were prepared by mixing 5μl of TaqMan probe, 18 μl of each primer, and 59μl of ddH 2O. Glyceraldehyde-3-phospha tedehydrogenase (GAPDH) and hypoxanthine-guanine phsphoribosyl-transferase (HPRT1) TaqMan assays (Fisher Scientific) were used as housekeeping gene controls. qRT-PCR was performed in 384-well plates, using dTTP MasterMix (Eurogentec), according to the manufacturer ’s instructions (for each well, 0.5 μl of TaqMan assay, 5μl of Mastermix , and 4.5 μl of cDNA). Amplification was performed as recommended by the manufacturer, on Roche LightCycler 480, and the relative quantification tool from the analysis software was applied. Data were analyzed using Microsoft Excel to calculate ΔC t and further analyzed in GraphPadPrism.

### In vitro T cell culture

Secondary lymphoid organs (mesenteric, axillary, inguinal, brachial, cervical, and lumbar lymph nodes and spleen) from SUV39H1-KO or WT OT-I male mice were collected, pooled and homogenized into single cell suspensions by scratching in a 40 µm cell strainer (Falcon) using a syringe plunge. Naïve (CD44-) CD8^+^ T cells were enriched using the naïve CD8_+_ T-cell isolation kit (Miltenyi 130-096-543) according to manufacturer’s instructions. Cells were then stained with Cell Trace Violet (Thermo Fisher). In brief cells were diluted at 10×10^6^ cells par ml of a solution of 5 µM of Cell Trace Violet in PBS during 15min 37C and then washed with complete medium [RPMI, 100 mM Hepes, 2 mM Glutamax, 50 uM 2-mercaptoethanol, 100 U/100 ug/ml Penicillin/Streptomycin, 1x MEM Non-essential amino acids, 1 mM Sodium Pyruvate (all Gibco, Thermo Fisher), 10% SBF (Eurobio). 1×10^4^ or 1×10^5^ CTV-stained naïve OTI CD8+ T cells were incubated for different time points with 2.5×10^4^ or 2.5×10^5^ splenocytes from CD3-KO mice preloaded with the indicated concentration of peptides (SIINFEKL, SIITFEKL or SIIVFEKL, GeneCust) in complete medium for the indicated periods of time. Then cells were stained for surface antibodies and analysed by FACS.

### Tumours

1 × 10^6^ B16-OVA-Luc cells were injected intravenously in the lateral tale vein of the mice. Tumor burden was measured by bioluminescence imaging using Xenogen IVIS Imaging System (PerkinElmer) two times per week. Acquired bioluminescence data was analyzed using Living Image software (PerkinElmer) and expressed in average radiance (photons/sec/cm^2^/sr). The humane endpoint was defined by a maximum of 20% weight loss and additional clinical signs of distress.

### CITE-seq library preparation and sequencing

SUV39H1 KO (CD45.1/1) and WT (CD45.1/2) OTI cells were adoptively transferred into Rag KO mice and 34 days after transfer mice were stained with CD45.1-PE antibody intravenously and euthanized 5’ after injection as indicated above. Enriched lung leucocytes were stained with the following antibodies: CD8α-PerCP, TCRb-PECy7, CD4-FITC, CD45.2-APC, B220-eF450, MHCII-eF450, Nkp46-V450 and DAPI. The following cell populations were sorted in a FACSAria FACS sorter: WT iv+ (CD45.1+/CD45.2+), WT iv- (CD45.1-/CD45.2+), KO iv+ (CD45.1+, CD45.2-), KO iv- (CD45.1-, CD45.2-) from 6 mice and pooled by population. After sorting, cells were stained with TotalSeq-A0078 anti-mouse CD49d, TotalSeq-A0112 anti-mouse CD62L, TotalSeq-A0073 anti-mouse/human CD44, TotalSeq-A0197 anti-mouse CD69, TotalSeq-A0013 anti-mouse Ly-6C Antibody, and each sorted population was stained with a different hashtag antibody: KOiv+, TotalSeq-A0301 anti-mouse Hashtag 1; WT iv+, TotalSeq-A0302 anti-mouse Hashtag 2; WT iv-, TotalSeq-A0303 anti-mouse Hashtag 3, KO iv-: TotalSeq-A0304 anti-mouse Hashtag 4 (all from Biolegend). Equal number of cells from each population were pooled and then subjected to 10X single cell RNA seq library preparation targeting 1×10^4^ total cells. The single-cell library method used was the Chromium Single Cell v3 with single-indexing. The RNA samples and antibody derived tag (ADT/HTO) samples were pooled separately. RNA samples were sequenced to a depth of 100,000 reads/cell on a NovaSeq S4.

### RNA-seq and TCRb library preparation and sequencing

For RNA and TCRb sequencing from ex-vivo naïve, memory and Trm populations, secondary lymphoid organs (mesenteric, axillary, inguinal, brachial, cervical and lumbar lymph nodes and spleen) from Suv39H1-KO or WT male mice were collected, pooled, homogenized into single cell suspensions and stained with CD4-V500, CD44-PerCPCy5.5, TCRb-FITC, CD122-PE, CD8a-APC-Cy7 and DAPI. Liver leucocytes were enriched as described before and stained with CD4-V500 CD69-PerCPCy5.5, Ly6c-AF488, TCRb-PECy7, CD8-APCCy7 and DAPI. Naïve (DAPI-/CD8a+/TCRb+/CD4-/CD44-/CD122-), and memory (DAPI-/CD8a+/TCRb+/CD4-/CD44+/CD122+) CD8^+^ T cells from pooled spleen and LN and CD69+ Trm (DAPI-/CD8a+/TCRb+/CD4-/CD69+/Ly6c-) from liver were FACS sorted directly into buffer RLT (Qiagen) supplemented with 2-mercaptoethanol (Sigma-Aldrich) and frozen at -80 C. Total RNA was isolated using RNeasy Plus Micro kit (Qiagen 74004) according to manufacturer’s instructions and quantified by NanoDrop (Thermo-Fisher). Total RNA size distribution was determined by RNA 6000 Pico Kit for 2100 Bioanalyzer systems (Agilent).

For RNA sequencing, cDNA library was prepared using a SMART-Seq v4 Ultra Low Input RNA kit (Takara) according to manufacturer’s instructions. Libraries were quantified by Qubit fluorometric quantification (Invitrogen) with dsDNA HS (High Sensitivity) Assay Kit and size distribution was determined by Agilent High Sensitivity DNA Kit. 100bp paired-end sequencing was performed on an Illumina NovaSeq – S1.

For TCRb sequencing, libraries were prepared from purified total RNA using iRepertoire kit for TCRb mouse sequencing according to manufacturer’s instructions. 150bp paired-end sequencing was performed on an Illumina MiSeq - nanoV2-300 with 10% PhiX.

For mRNA sequencing from in vitro activated cells, cells were stained with DAPI, CD8a-PE and TCRb5 APC and DAPI-/CD8a+/TCRb+ cells were FACS sorted in a FACSAria Fusion directly into buffer RLT (Qiagen) supplemented with 2-mercaptoethanol (Sigma-Aldrich) and frozen at -80 C. Total RNA was isolated using RNeasy Plus Micro kit (Qiagen 74004) according to the manufacturer’s instructions. RNA was quantified by NanoDrop (Thermo-Fisher) and size distribution was analyzed by using Agilent RNA 6000 Nano kit (Agilent).

mRNA sequencing libraries were prepared from 100ng of total RNA using the Stranded mRNA Prep Ligation library preparation kit (Illumina), which allows to perform a strand specific sequencing. This protocol includes a first step of polyA selection using magnetic beads to focus sequencing on polyadenylated transcripts. After RNA fragmentation, cDNA synthesis was then performed and resulting fragments were used for dA-tailing followed by ligation of RNA Index Anchors. PCR amplification with indexed primers (IDT for Illumina RNA UD Indexes) was finally achieved, with 15 cycles, to generate the final cDNA libraries.

Individual library quantification and quality assessment was performed using Qubit fluorometric assay (Invitrogen) with dsDNA HS (High Sensitivity) Assay Kit and LabChip GX Touch using a High Sensitivity DNA chip (Perkin Elmer). Libraries were then equimolarly pooled and quantified by qPCR using the KAPA library quantification kit (Roche). Sequencing was carried out on the NovaSeq 6000 instrument from Illumina using paired-end 2 × 100 bp, to obtain around 35 million clusters (70 million raw paired-end reads) per sample.

### Bulk RNA-seq analysis

Bulk RNA-seq fastq reads were mapped to reference mouse genome mm10 using STAR (v2.7.1.a). Gene expression was quantified using featureCounts from Subread (v1.6.4) with Gencode mouse genes annotations (gtf file release M22, GRCm38.p6) and default parameters. Genes raw counts matrices were imported to R (v4.2.1). TPM and FPKM normalized expression values were calculated using calculateFPKM and calculateTPM functions from scuttle R package (version 1.8.4). Low expressed genes were filtered out using a threshold of 0.5 FPKM. Only genes expressed above this threshold in all samples from at least 1 condition were retained. Count normalization and differential expression analyses were performed using DESeq2 R package (version 1.36.0). Genes with Benjamin-Hochberg (BH) adjusted p-value below 0.05 were considered as significantly differentially expressed between conditions. Gene Set Enrichment Analyses (GSEA) were performed using GSEA software (v4.3.2) from MsigDB with curated gene signatures and default parameters: Number of permutations = 1000; Permutation type = phenotype when more than 7 samples were compared in each condition otherwise gene_set). Bubble gum representations were created within R using R package ggplot2 (v3.4.4).

### CITE-seq analysis

Single cell RNA-seq fastq reads for gene expression were processed using the Cell Ranger Single Cell Software Suite (v3.1.0) to perform quality control, barcode and UMI processing, and single cell 3ʹ gene counting. Reads were mapped to mm10 transcriptome (v3.0.0). HTO (hashtag oligos) sequencing reads defining sample of origin and ADT (antibody-derived tags) sequencing reads assessing cell surface protein expression were processed using CITE-seq count (v1.4.5). Single cell gene expression data were imported within R (v4.2.1) and downstream analyses were performed using Seurat R package (v4.3.0.1). Cells with less than 200 features expressed and/or more than 10% of mitochondrial content were filtered out. ADT and HTO processed data from CITE-seq count were then imported in the Seurat object in two new assays. After CLR normalization, HTO data were demultiplexed with Seurat function HTODemux to assess sample of origin of each cell (blood-KO, blood-WT, tissue-KO, tissue-WT). Only singlet cells were kept for downstream analysis while cells positive for two or more hashtag oligos (doublets) and cells negative for all hashtag oligos were removed. For protein level assessment, counts from the ADT assay were normalized using the CLR method and used in downstream analyses to support annotation of the clusters defined by gene expression. For gene expression analysis, after log-normalization, the 5000 most variable genes were scaled and used to compute Principal Component Analysis (PCA). Top 30 principal components were chosen to compute UMAP and performed clustering steps. Differential expression analysis on gene expression was computed on clusters identified at resolutions 0.8 and 1.2. Scores for curated signatures were computed using AddModuleScore function from Seurat. Prediction of differentiation state of cells was computed using R package CytoTRACE (v0.3.3). Transcription factor and regulon activities were evaluated using R package SCENIC (v1.3.1). Trajectory analysis was performed using R tools suite dynverse (dyno v0.1.2, dynmethods v1.0.5, dynwrap 1.2.4, dynguidelines v1.0.1, dynfeature 1.0.1, dynplot 1.1.2, dynutils 1.0.11, dynparam 1.0.2, dyndimred 1.0.4). Differentially expressed genes excluding genes from the cell cycle cluster were used for the input matrix in the trajectory analysis. Following recommendations from dynguidelines::guidelines function on the dataset, slingshot (ti_slingshot) was used as the trajectory inference method within infer_trajectory function from dynwrap. Abundance of spliced and unspliced reads was evaluated using velocyto (v0.17.17) on cellranger output. From velocyto loom file output, python package scvelo (v0.2.5) was used to process, assess, and represent RNA velocity. Silhouette scores for each cluster were computed using the silhouette function from R package cluster (v2.1.4). For computing original proportions of each population (WTiv+, KOiv+, WTiv- and KOiv-) within each cluster we used formula 1:

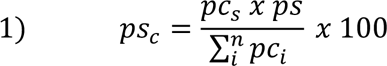

Where ps_c_ is the proportion of sample s in cluster c, pc_s_ is the proportion of cluster c in sample s, ps is the proportion of sample s among the whole lung and pc_i_ is the proportion of cluster c in sample i.

### Statistical analysis and plots

Heatmaps, barplots, violin plots, dotplots, scatter plots, UMAP representations and volcano plots of single cell and bulk sequencing data were computed within R (v4.2.1) using R packages ggplot2 (v3.4.4), Seurat (v4.3.0.1), ComplexHeatmap (v2.14.0), scvelo (v0.2.5) and dynplot (v1.1.2). Statistical analysis and other plots were made by GraphPad Prism version 10.11. Plots were exported into Affinity Designer version 1.10.8 to build the figures.

