## Supplementary material for "Epigenetic control of CD8^+^ T cell tissue homing and tissue resident memory T cell precursors by the histone methyltransferase SUV39H1": Supp figures

**Supplementary Figure 1: SUV39H1-deficiency leads to the accumulation of CD8⁺ tissue-homing CD8⁺ T cells at steady-state**

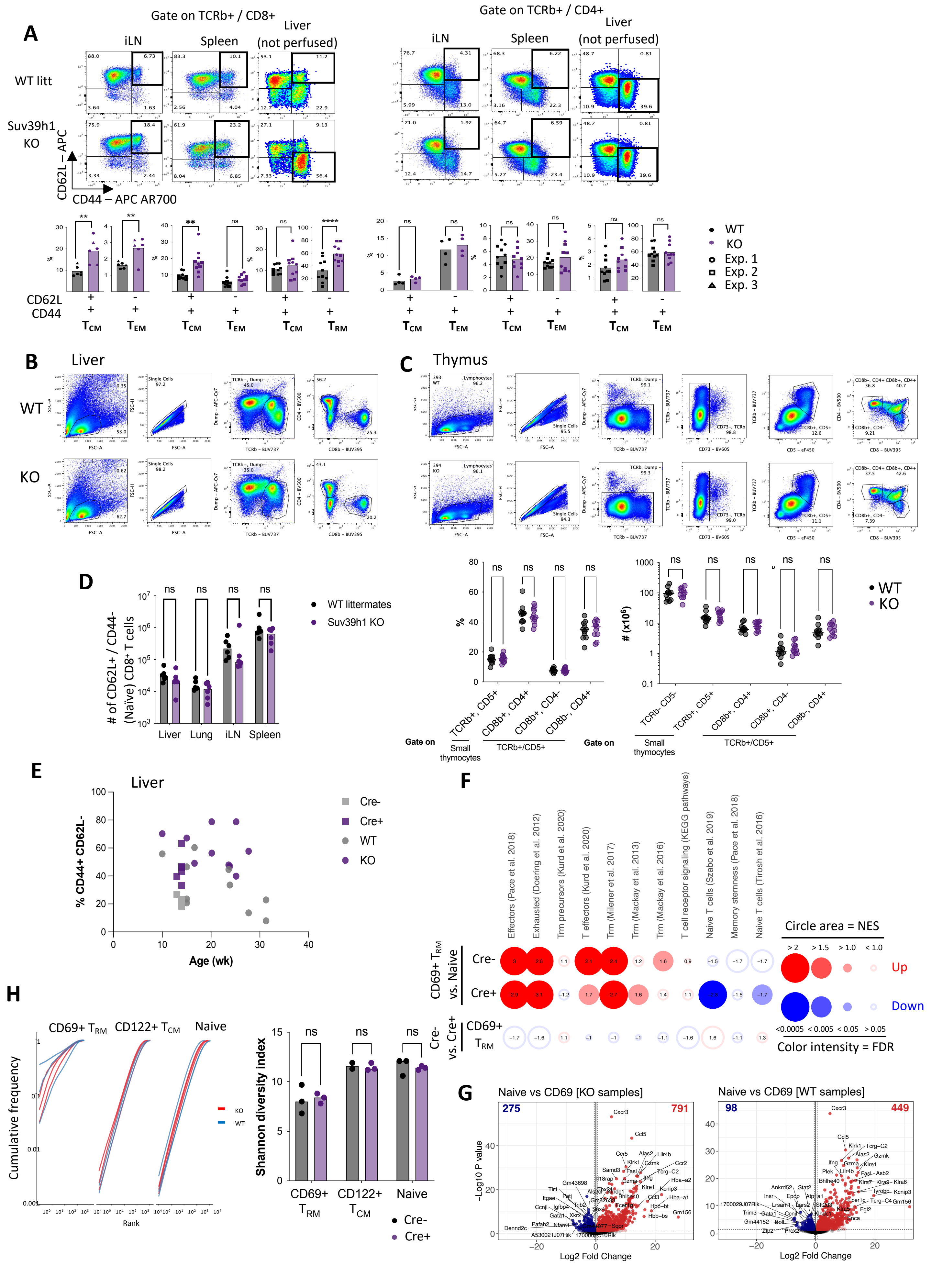

**Supplementary Figure 1: SUV39H1-deficiency leads to the accumulation of $CD8^+$ tissue-homing $CD8^+$ T cells at steady-state**

**A.** Representative dot plots and quantification of CD44 and CD62L expression in $CD8^+$ (**left**) or $CD4^+$ (**right**) T cells from the indicated organs of full SUV39H1 KO or WT littermates at steady state.

**B.** Gating strategy for the analysis of data shown in Figure 1. Figure shows two liver samples as examples.

**C**. Representative gating strategy, plots and quantification of the indicated populations from thymus samples from the same mice as in Figure 1 A.

**D.** Number of naïve (CD44- CD62L+) CD8+ T cells in the indicated organs from the same mice as in Figure 1 A.

**E.** Plot of the percentage of $T_{RM}$ (CD44+ CD62L-) CD8+ T cells in liver according to the age of the mice for mice from Figure 1 A and B.

**F.** GSEA analysis of RNAseq data from CD69+ $T_{RM}$ CD8+ T cells from liver vs. Naïve CD8+ T cells from pooled LN and spleen from conditional SUV39H1 mice (Cre+) or WT littermates (Cre-) like in (E**)** or CD69+ $T_{RM}$ from WT littermates (Cre-) vs. CD69+ $T_{RM}$ from conditional SUV39H1 KO (Cre+) mice.

**G**. Volcano plots of the comparison between CD69+ $T_{RM}$ CD8+ T cells from liver vs. Naïve CD8+ T cells from pooled LN and spleen from conditional SUV39H1 mice (Cre+) or WT littermates (Cre-) like in (E).

**H.** TRCb CDR sequencing analysis in $T_{RM}$ (CD69+ Ly6c- CD44+ CD62L-) CD8+ T cells from liver and $T_{CM}$ (CD122+ CD44+) and Naïve (CD122- CD44-) CD8+ T cells from pooled LN and spleen.

ns: not significant by multiple unpaired t test (B and C) and multiple paired t test (E).

**Supplementary Figure 2: CD8$^+$ T$_{RM}$ differentiation upon flu infection is enhanced in the absence of SUV39H1**

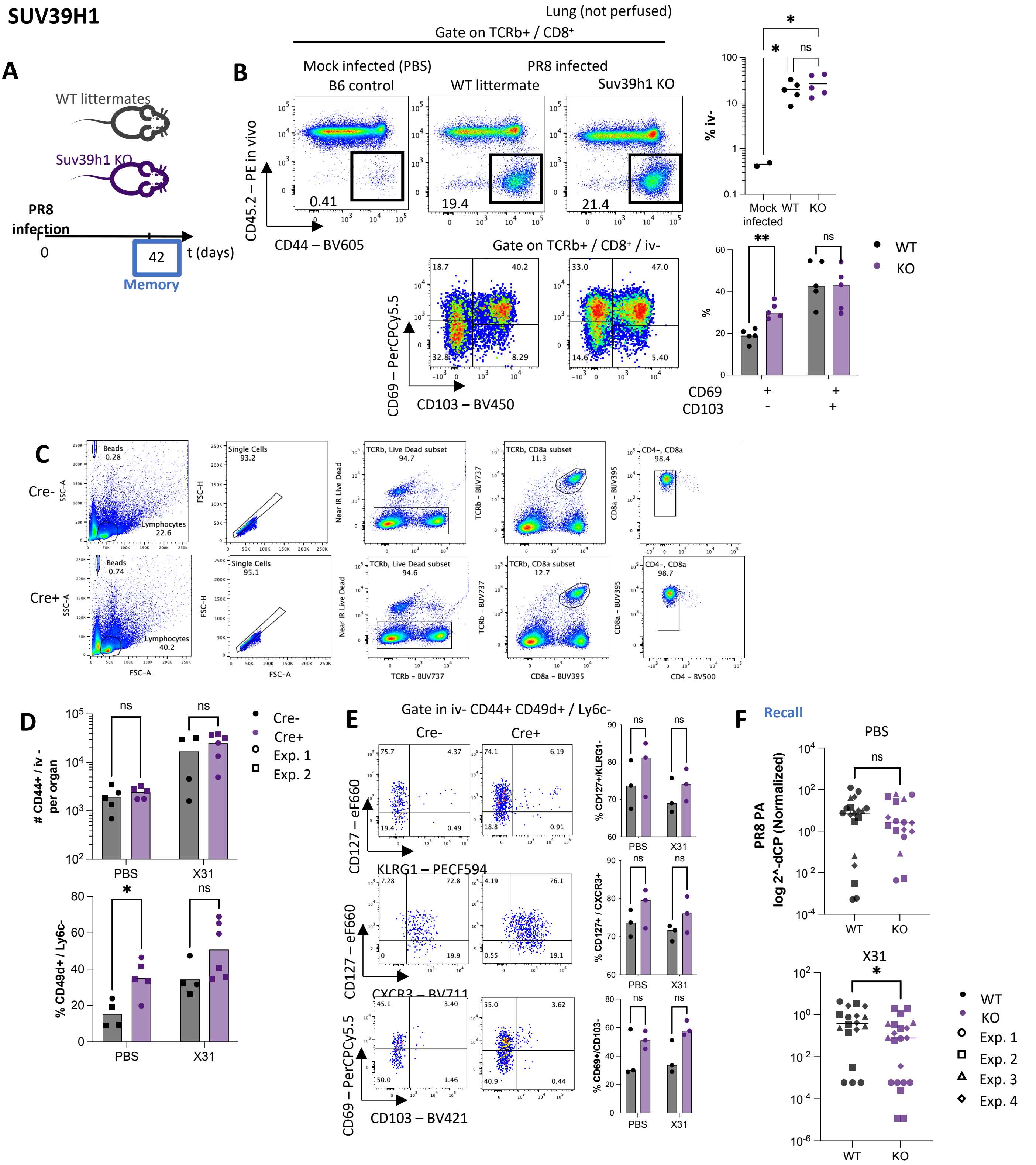


**Supplementary Figure 2: CD8$^+$ T$_{RM}$ differentiation upon flu infection is enhanced in the absence of SUV39H1**

**A.** Scheme of the experiment.

**B.** Representative dot plots and quantification of the % of iv- CD8$^+$ T cells (top) and CD69/CD103 expression in iv- CD8$^+$ T cells (bottom) in lungs of SUV39H1 KO or WT littermates 42 days after PR8 infection.

**C.** Gating strategy for the analysis of data shown in Figure 3 B and C.

**D.** Raw data used to build normalized data in Figure 3 B.

**E.** Representative dot plots and quantification of the expression of the indicated markers in CD44+ iv- CD49d+ Ly6c- CD8+ T cells from lungs during the memory phase as in Figure 3 B.

**F.** PR8 PA quantification by qPCR in lungs 3 days after PR8 infection (**Recall**) as in Figure 3.

* p<0.05 ; ns: not significant by ANOVA test with Šídák's multiple comparisons test (**D** and **E**) or Mann Whitney test (**F**).

### Supplementary figure 3. CITE-seq analysis of lung-homing CD8⁺ T cells

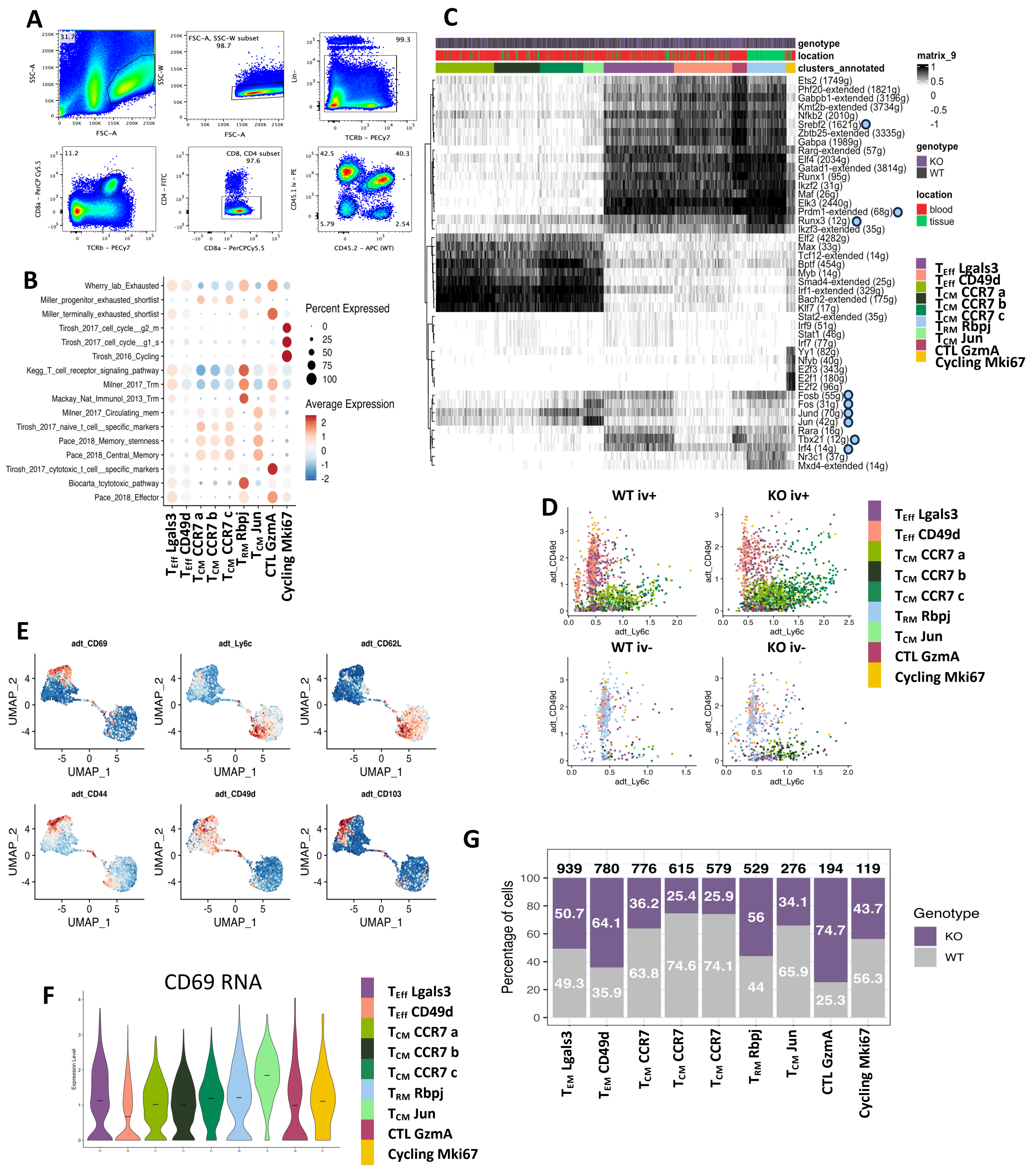

**Supplementary Figure 3. CITE-seq analysis of lung-homing CD8+ T cells**

**A.** Sorting strategy and original proportions of cells in lungs. Lin: MHCII, Nkp46

**B.** Dot plot showing the average expression of the indicated RNA signatures in each cluster at 0.8 resolution.

**C.** Heatmap representing binary activity for each cell of the top 10 transcription factors (regulons) with the highest foldchange in each cluster at 0.8 resolution. Transcription factor activity was assessed using SCENIC algorithm. Numbers indicate the number of genes for each regulon. Circles indicate TF known to be involved in $T_{RM}$ biology.

**D.** Dot plots showing the expression of CD49d and Ly6c proteins assessed through antibody tagging of the CITE-seq experiment. Each dot represents a cell. Cells are color-coded by clusters at 0.8 resolution.

**E.** Protein expression by antibody tagging of the 6 markers analysed in the CITE-seq experiment. Each dot represents a cell.

**F.** Volcano plot of Cd69 RNA expression for each cluster at 0.8 resolution.

**G.** Genotype distribution of cells for each cluster at 0.8 resolution.

### Supplementary figure 4. SUV39H1 regulates the developmental pathway of $T_{RM}$ cells

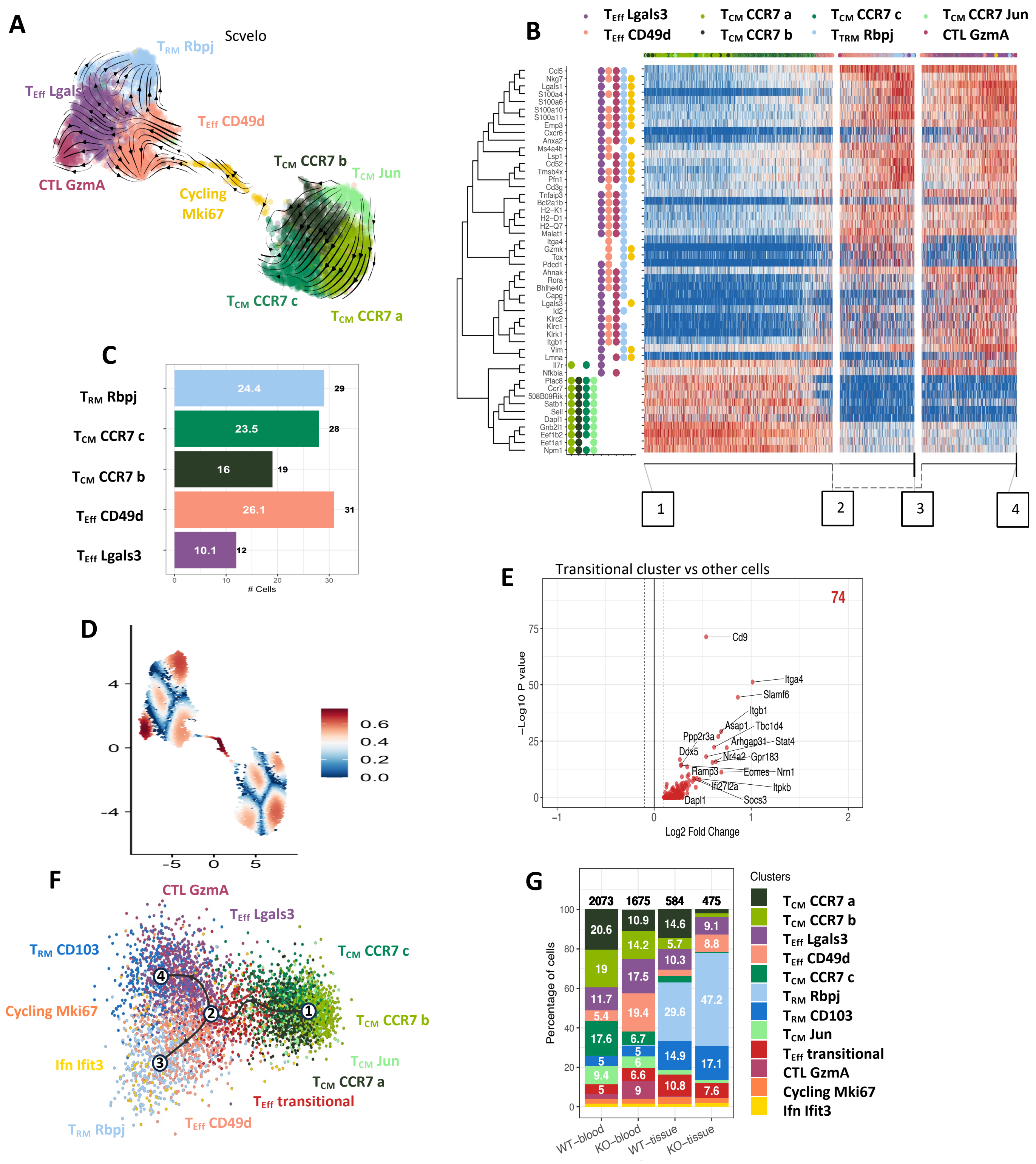


**Supplementary Figure 4. SUV39H1 regulates the developmental pathway of $T_{RM}$ cells.**

**A.** Scvelo trajectory analysis. Each dot represents a cell. Cells are color-coded by cluster at 0.8 resolution.

B. Heatmap expression of the top 50 genes varying the most along the trajectory ordered by Slingshot pseudotime. Squares indicate milestones. Coloured dots in the left indicate significant (q<0.05) expression of each gene in each cluster at 0.8 resolution by DE analysis.

**C.** Label transfer prediction distribution of cycling cells onto other clusters at 0.8 resolution. Numbers within the bars represent the percentage of the total cycling cluster and numbers over bars represent the total number of cells falling in each cluster.

**D.** UMAP representation of the single cell RNAseq data showing SIhouette scores at resolution 0.8.

**E.** Volcano plot showing genes differentially upregulated by transitional cluster vs all other cells.

**F.** Trajectory analysis results using Slingshot inference method. Cells are color-coded by clusters identified using resolution 1.2.

**G.** Distribution of clusters at 1.2 resolution in each sample. Numbers within bars indicate percentages of the total cells in the sample. Numbers over bar represent total number of cells for each sample.

### Supplementary Figure 5: $T_{RM}$ precursors formation is increased in the absence of SUV39H1

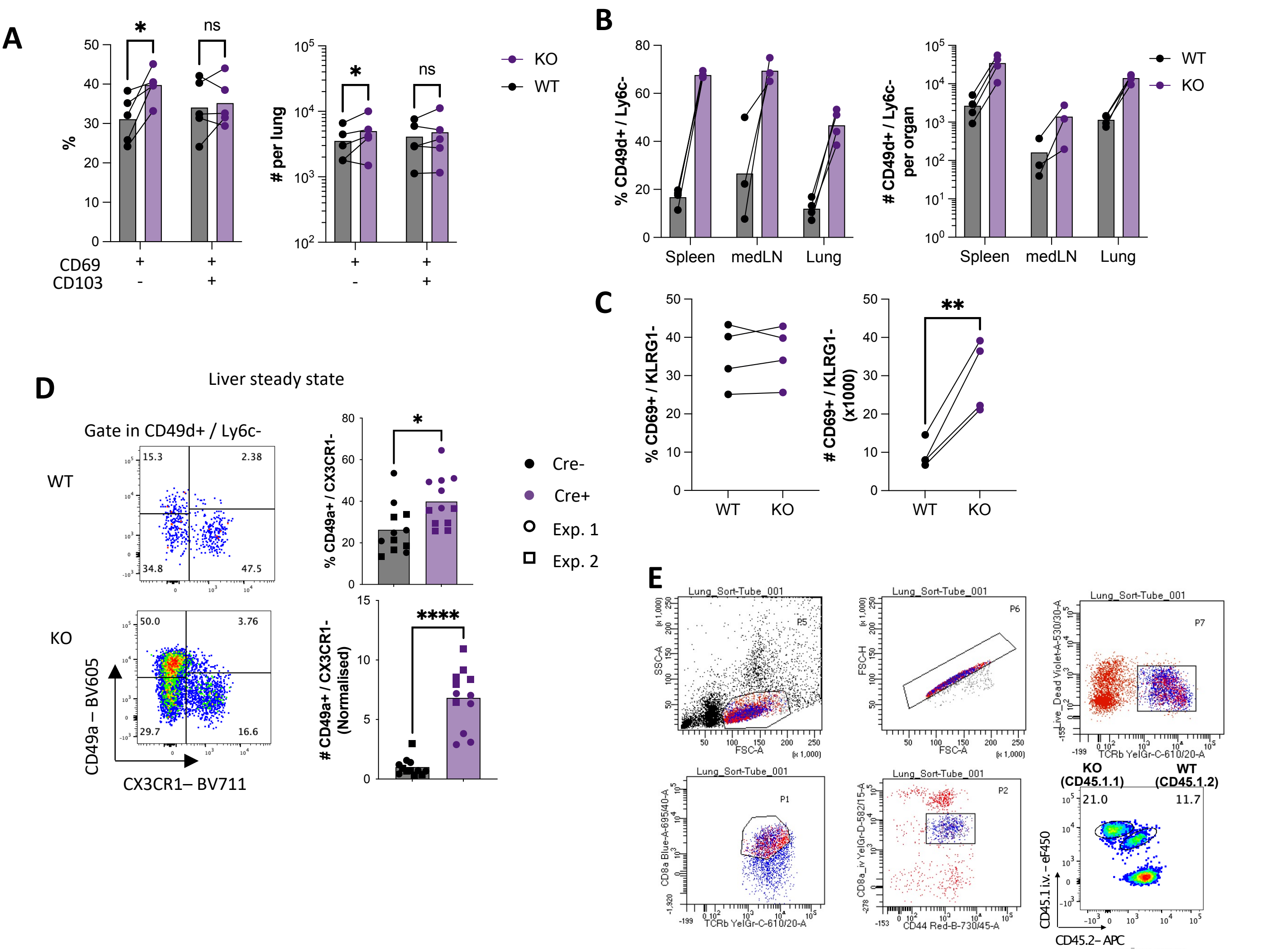


**Supplementary Figure 5: $T_{RM}$ precursors formation is increased in the absence of SUV39H1**

**A.** CD69/CD103 expression 30 days post infection in lungs (Memory) of replicate experiment from Figure 5 B.

**B.** Percentages and numbers of CD49d+/Ly6c- cells among OTI cells in the indicated organs 9 days post X31-OVA infection (Acute) of replicate experiment from Figure 6 C.

**C**. Percentages and numbers of CD69+/KLRG1- cells among CD49d+/Ly6c- cells in lungs 9 days post X31-OVA infection (Acute) of replicate experiment from Figure 6 C.

**D.** Representative dot plots and quantification of CD49a/CX3CR1 expression in CD44+ CD49d+ Ly6c- CD8$^{+}$ T cells from liver at steady state (as in Figure 1 D).

**E.** Sorting strategy of WT and SUV39H1-KO Total iv- CD8+ T cells from lungs from experiment in Figure 5 D and E.

p<0.05 ; ** p<0.01 ; **** p<0.0001 ; ns: not significant by multiple ratio paired t test (A) and (B), ratio pared t test (C) and unpaired t test (D).

### Supplementary Figure 6. SUV39H1-deficient lung-homing CD8+ T cells delay lung tumor metastasis growth

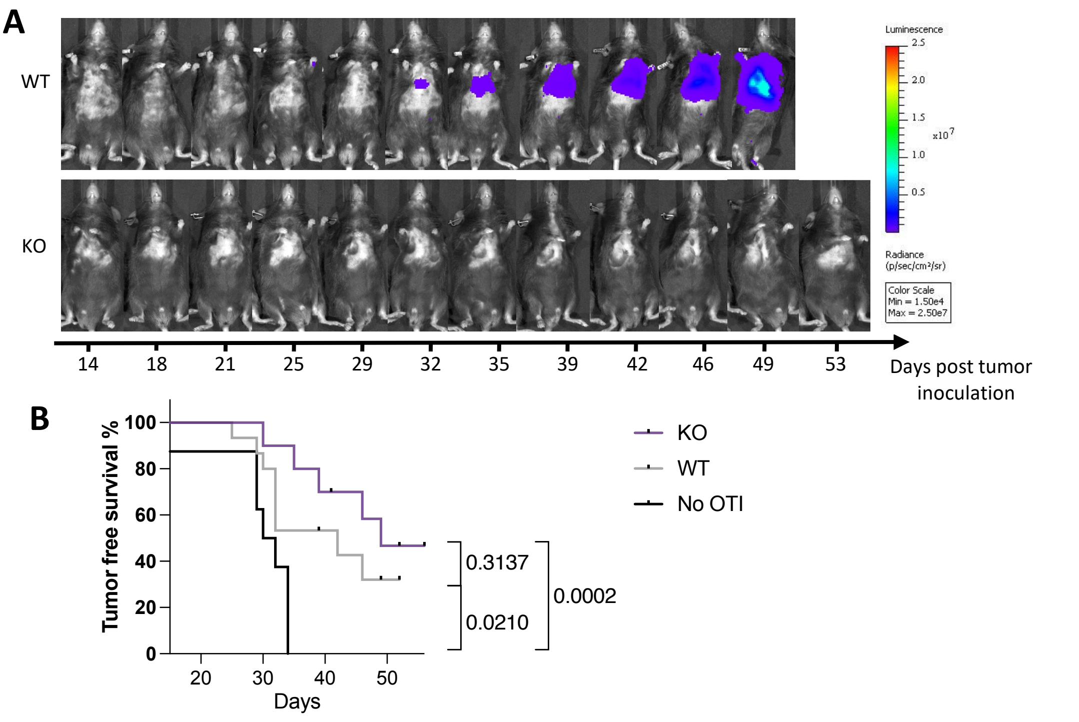


**Supplementary Figure 6. SUV39H1-deficient lung-homing CD8+ T cells delay lung tumor metastasis growth.**
**A.** Representative bioluminescent images of mice adoptively transferred with WT and KO OTI cells from Figure 6.
**B.** Tumor free survival curves (cut off average radiance = $10^3$ p/s/cm/sr, right) from Figure 6 C. Pooled of 2 independent experiments. Numbers represent p values by Log-rank (Mantel-Cox) test.

**Suplementary Table 1. $T_{RM}$ flow cytometry panel used in Figure 1 C.**

| Fluorochrome | Molecule |
|---|---|
| BUV737 | TCR |
| BUV395 | CD8b |
| eF450 | CD103 |
| V500 | CD4 |
| BV605 | CD49a |
| BV711 | CX3CR1 |
| BV786 | CD49d |
| PerCP Cy5.5 | CD69 |
| FITC | CD127 |
| PE | CD45.2 iv |
| PE-CF594 | CD122 |
| PE Cy7 | CD62L |
| APC | Ly6c |
| AR700 | CD44 |
